# Sample demultiplexing, multiplet detection, experiment planning and novel cell type verification in single cell sequencing

**DOI:** 10.1101/828483

**Authors:** Hongyi Xin, Qi Yan, Yale Jiang, Qiuyu Lian, Jiadi Luo, Carla Erb, Richard Duerr, Kong Chen, Wei Chen

**Author notes:** Corresponding authors. Contact: W.C.,. K.C.,.

## Abstract

Identifying and removing multiplets from downstream analysis is essential to improve the scalability and reliability of single cell RNA sequencing (scRNA-seq). High multiplet rates create artificial cell types in the dataset. Sample barcoding, including the cell hashing technology and the MULTI-seq technology, enables analytical identification of a fraction of multiplets in a scRNA-seq dataset.

We propose a Gaussian-mixture-model-based multiplet identification method, GMM-Demux. GMM-Demux accurately identifies and removes the sample-barcoding-detectable multiplets and estimates the percentage of sample-barcoding-undetectable multiplets in the remaining dataset. GMM-Demux describes the droplet formation process with an augmented binomial probabilistic model, and uses the model to authenticate cell types discovered from a scRNA-seq dataset.

We conducted two cell-hashing experiments, collected a public cell-hashing dataset, and generated a simulated cell-hashing dataset. We compared the classification result of GMM-Demux against a state-of-the-art heuristic-based classifier. We show that GMM-Demux is more accurate, more stable, reduces the error rate by up to 69×, and is capable of reliably recognizing 9 multiplet-induced fake cell types and 8 real cell types in a PBMC scRNA-seq dataset.

## 1. Introduction

Droplet-based single cell RNA sequencing (scRNA-seq)[18, 13, 45] has provided many valuable insights into complex biological systems, such as rare cell type identification[41, 39, 25, 32], differential expression analysis at the single cell level[8, 5, 2], and cell lineage studies[8, 29, 15, 23]. While the per-cell cost of library prep has decreased over the years, the scalability of droplet-based scRNA-seq remains limited, mostly due to rapidly increasing, yet hard to anticipate, multiplet rates as more cells are loaded during single sequencing cell library prep[17]. Multiplets significantly confound the analysis of single-cell experiments and can lead to false discoveries[9, 17], such as false lineages in cell lineage tracing[14, 28, 20], incorrect categorizations in cell type classification[46, 43, 26], or false findings in rare cell type discovery[44, 22]. Large cell populations are especially required for rare cell type discovery, but loading large cell populations during scRNA-seq library prep leads to high multiplet rates. As a result, researchers are challenged with identifying real rare-type cells in a multiplet-filled scRNA-seq dataset. Overall, the scalability of scRNA-seq can be significantly improved, greatly reducing the per-cell library prep cost, if multiplets can be identified and removed from downstream analysis. To achieve greater adoption of single cell sequencing technology, it is crucial to 1) identify and remove multiplets from downstream analysis, 2) anticipate the multiplet rate prior to conducting an experiment, and 3) verify whether rare cell types identified from a single cell dataset are authentic and are not multiplets.

Recently, emerging sample barcoding technologies, such as cell hashing[37] or MULTI-seq[21], enable identification of multiplets arising from more than one uniquely labeled sample and their subsequent removal from downstream analysis of single cell sequencing experiments. Both methods use oligonucleotide-labeled reagents that conjugate on the cell surface to produce sample-specific markings on cells: cell hashing, an extension of the cellular indexing of transcriptomes and epitopes by sequencing (CITE-seq) technology[36], uses barcoded oligo-conjugated antibodies that targets ubiquitously expressed surface markers, such as CD298 and beta2-microglobulin, while MULTI-seq uses lipid- and cholesterol-modified oligonucleotides that attach to the cell surface membrane and the cell nuclei membrane. For simplicity, we refer to the oligonucleotide-labeled reagents used in both methods as sample-hashtag oligonucleotides (HTO). Sample barcoding involves labeling cells from each sample with sample-specific HTO conjugates and then pooling the HTO-labeled cells from different samples for droplet-based scRNA-seq sequencing library prep. During library prep, the pooled cell assay is driven through a microfluidic chip to form cell-assay droplets. A fraction of cell-assay droplets are combined with barcode-enclosing gel beads and form Gel Beads in Emulsion, or GEMs. Inside each GEM, HTO barcodes are combined with GEM barcodes. Subsequent sequencing simultaneously recovers the HTO barcode(s) and the GEM barcode for each GEM. An abstract workflow of a 3-sample sample barcoding experiment is provided in Figure 1. Finally, the count of the HTO unique molecular identifiers (UMIs) for each sample, which translates to the number of cell-attached, sample-specific HTO antibodies of each GEM are summarized in a matrix, called the HTO matrix. Table 1 depicts an example 3-sample HTO matrix.

**Figure 1:**
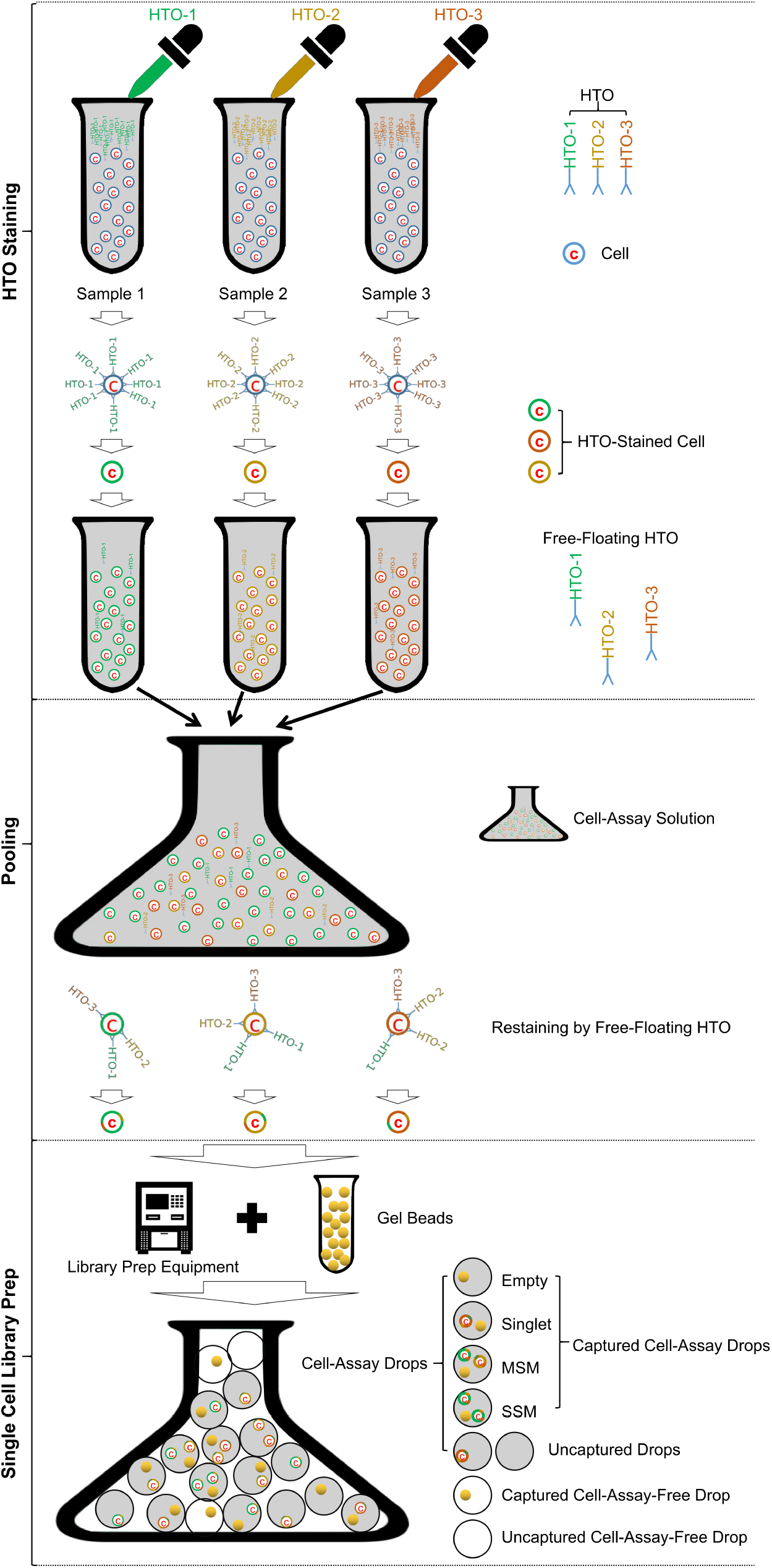
Sample barcoding workflow.

**Table 1:**
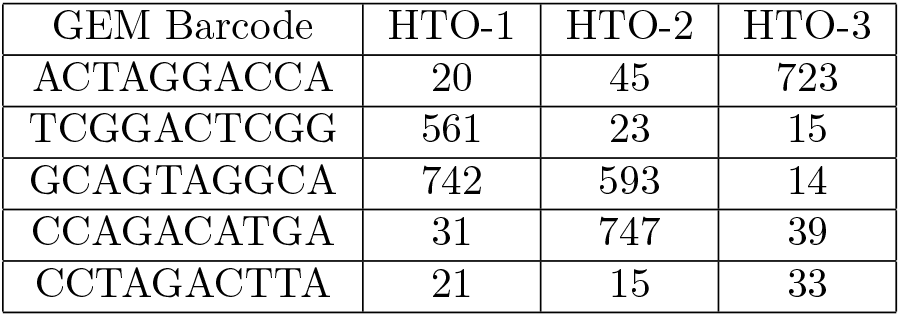
An example HTO matrix. Each row is a GEM with its unique GEM-barcode as index. Each column is a HTO sample id. The *i*th row and *j*th column of the matrix stores the number of HTO antibodies (in the form of UMI counts) of the *j*th HTO sample (HTO-*j*) attached to cells in the *i*th GEM.

There are three types of droplets in a sample barcoding scRNA-seq dataset: 1) *Multi-sample multiplets (MSMs):* droplets that contain more than one cells from more than one HTO samples; 2) *Single-sample multiplets (SSMs):* droplets that contain more than one cells from the same HTO sample. 3) *Singlets*: droplets that contain a single cell. We combine singlets and SSMs into a single category called *single-sample droplets (SSDs)* to differentiate them from MSMs. The relationship between MSM, SSM, singlet and SSD are summarized in Figure 2. MSMs can be distinguished from SSDs in the HTO matrix: MSMs typically have high HTO UMI counts for more than one HTO barcode, while SSDs typically have high UMI counts for only one HTO barcode and low HTO UMI counts for all other HTO barcodes. However, sample barcoding cannot separate singlets from SSMs, as these two droplet types are indistinguishable in the HTO matrix. As a result, SSMs cannot be removed by sample barcoding and will remain in the dataset as noise.

**Figure 2:**
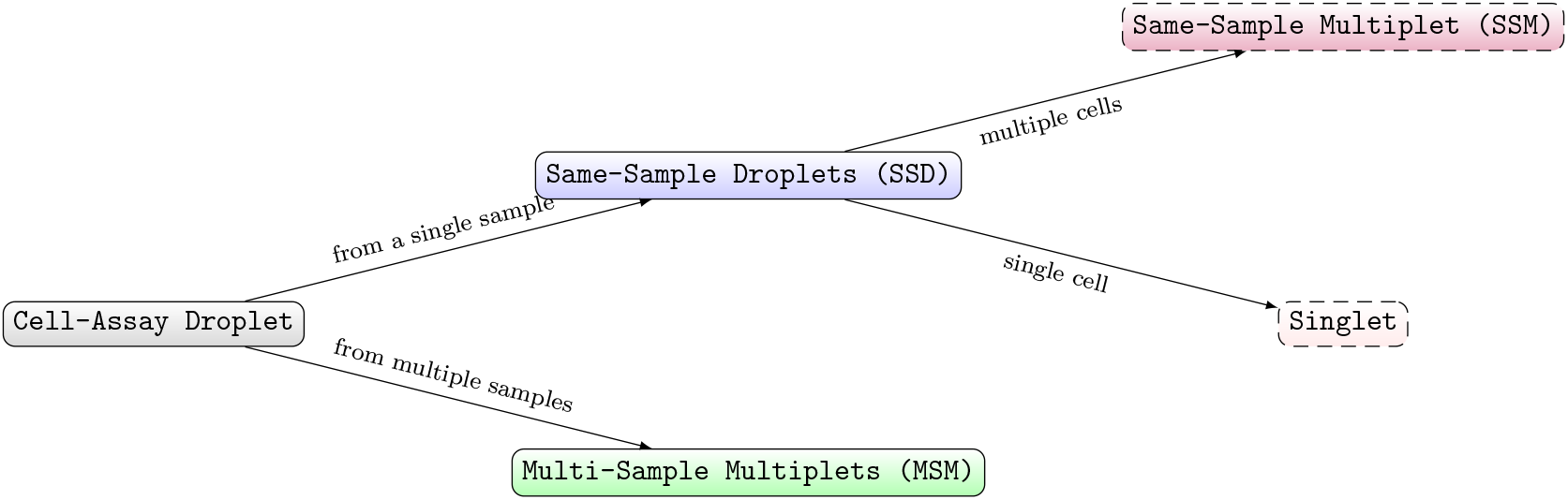
Relationship between MSM, SSM, SSD and singlet. SSD and MSM are differentiated based on whether the droplet contains cells from multiple HTO samples. SSM and singlet are further differentiated by the number of cells in the droplet.

GEMs can also be classified based on the number of cell types enclosed in them. GEMs that contain a single cell type are named *pure-type GEM*whereas GEMs that contain multiple cell types are named *phony-type GEMs*. An illustration of phony-type GEMs and pure-type GEMs is provided in Figure 3A. Pure-type GEMs are not necessarily singlets—a pure-type GEM can still be a multiplet, but contains cells of exactly the same cell type. Hence a pure-type GEM could be a singlet, a MSM or a SSM. Phony-type GEMs, on the other hand, are all multiplets. Hence they must be either MSMs or SSMs.

**Figure 3:**
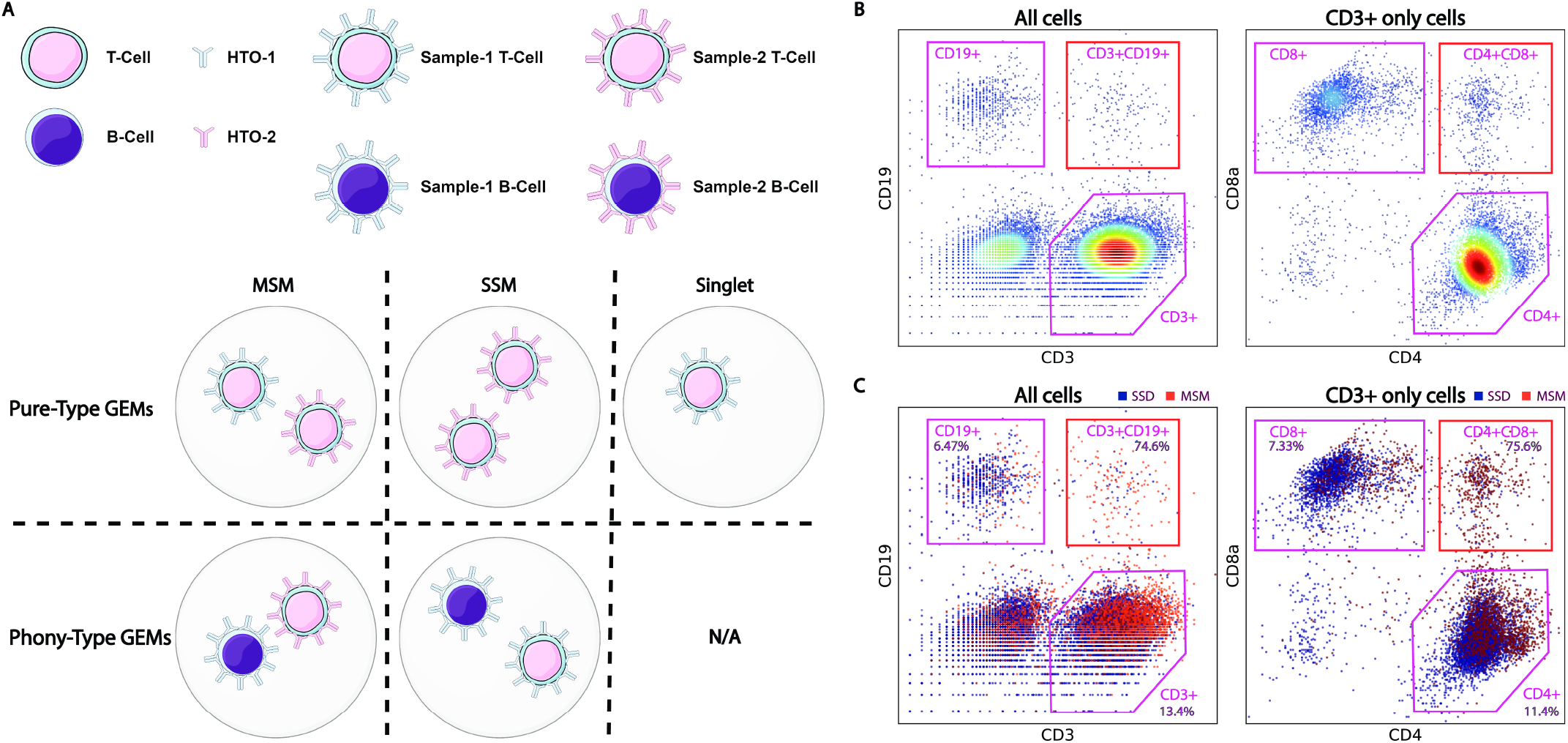
Examples of pure-type and phony-type GEMs in scRNA-seq data. Fig. A shows example compositions of pure-type and phony-type GEMs in a cell hashing dataset. Note that phony-type GEMs cannot be singlets. Fig. B presents the gating results of a 4-sample cell hashing CITE-seq PBMC dataset. Notice the unconventional cell types in B (highlighted in red bounding boxes). Fig. C shows the MSM ratio of each cell type in B. The alleged novel cell types in B are all phony-type cells, highlighted by their high MSM ratios.

Phony-type GEMs can be misclassified as novel rare cell types. Figure 3B depicts the gating results of a 16K-GEM PBMC cell hashing and CITE-seq dataset. The CD3-CD19 scatter plot shows GEMs comprised of CD19^+^ B cells, CD3^+^ T cells, and also a CD3^+^CD19^+^ double positive T-B cell GEM cluster. Similarly, in the CD4-CD8 scatter plot of all CD3^+^ T cells, besides the CD4^+^ helper T cells and CD8^+^ cytotoxic T cells, there also exists a CD4^+^CD8^+^ helper-cytotoxic T cell GEM cluster. Both clusters are highlighted in red circles. Existence of such PBMC types defies conventional wisdom, where CD3^+^CD19^+^ cells and CD4^+^CD8^+^ T cells are believed to be extremely rare[1, 33]. In fact, as revealed by cell hashing, both cell types, along with many other alleged novel rare cell types discovered in this dataset, are all phony cell types. Most, if not all, GEMs in these phony-cell-type clusters are phony GEMs: instead of containing real T-B cell(s), each CD3^+^CD19^+^ GEM is amultiplet that contains individual CD3^+^ T and CD19^+^B cell(s). When compared against true cell types, such as CD19^+^ B cells or CD4^+^ helper T cells, phony-type GEMs are most likely to be MSMs. Figure3C displays the MSM ratios of the CD19^+^ B cell, the CD4^+^ helper T cell and the CD8^+^ cytotoxic T cell true-cell-type GEM clusters (also referred to as pure-type GEM clusters); as well as the MSM ratios of the CD3^+^CD19^+^ and the CD4^+^CD8^+^ phonycell-type GEM clusters (or simply phony-type GEM clusters). From the figure we observe that phonycell-type GEM clusters have much higher MSM ratios than true-cell-type GEM clusters (~ 75% vs. < 14%).

The existing MSM classifier[37] provided by the single cell analysis toolkit Seurat[4], referred to as the *heuristic classifier* in this paper, for processing sample barcoding data is heuristic-based and unstable: it generates inconsistent results in repetitive runs and generates diverging results under different heuristic parameters. It does not model SSM. Therefore, it cannot estimate the percentage of singlets and SSMs in the dataset; and it cannot predict the percentages of MSMs, singlets and SSMs of the conceived output of a planned sample barcoding experiment. Most importantly, it cannot determine whether an alleged novel-cell-type-defining GEM cluster consists of mainly pure-type GEMs. Hence it cannot use sample barcoding results to verify the legitimacy of putative novel cell types in scRNA-seq.

In this work, we propose a model-based Bayesian framework, GMM-Demux, for sample barcoding data processing. GMM-Demux consistently and accurately separates MSMs from SSDs; estimates the percentage of SSMs and singlets among SSDs; anticipates the MSM, SSM and singlet rates of planned future sample barcoding experiments; and verifies the legitimacy of putative novel cell types discovered in sample-barcoded scRNA-seq datasets. Specifically, GMM-Demux independently fits the HTO UMI counts of each sample into a Gaussian-mixture model[34]. From each Gaussian-mixture model, GMM-Demux computes the posterior probability of a GEM containing cells from the corresponding sample. From the posterior probabilities, GMM-Demux computes the probabilities of a GEM being a MSM or a SSD. Among SSDs, GMM-Demux estimates the proportion of SSMs and singlets in each sample using an augmented binomial probabilistic model. Using the probabilistic model, GMM-Demux checks if a proposed cell-type defining GEM cluster is a pure-type GEM cluster or a phony-type GEM cluster; and based on the classification of the GEM cluster, GMM-Demux proves or disproves the novel-cell-type proposition.

To benchmark the performance of GMM-Demux, we conducted two in-house cell hashing and CITE-seq experiments, collected a public cell hashing dataset, and simulated an in-silico cell hashing dataset. We compare GMM-Demux against a human-supervised classifier and an existing, state-of-the-art heuristic classifier and show that GMM-Demux is the most accurate and the most consistent classifier, which reduces the classification error rate by up to 69×. Based on the probabilistic model of GMM-Demux, we build a publicly-accessible online sample-barcoded single-cell experiment planner that takes experiment parameters, including the planned number of cells and the planned number of HTO samples as inputs, and estimates the MSM, SSM and singlet rates of the anticipated sequencing outcome. We show that our online planner is compatible with both cell hashing experiments and ordinary droplet-based single cell sequencing experiments (which are equivalent to cell hashing experiments using a single HTO sample). Using the online experiment planner, we profile the multiplet rates over an array of experiment parameters and show that the estimated multiplet rates are consistent with observations from previous studies. From the cell hashing and CITE-seq PBMC dataset, we extracted 17 cell-type-defining GEM clusters through in-silico gating, based on surface marker expression levels. Among them, 9 GEM clusters define novel cell types that were previously thought to be non-existent in PBMC. Further analysis by GMM-Demux shows that all 9 novel-cell-type-defining GEM clusters are indeed phony-type GEM clusters and are removed from the dataset. The remaining 8 GEM clusters, on the other hand, define real cell types are consistent with existing knowledge. With GMM-Demux, we were able to load 35.7K cells during a single library prep and recover 15.8K GEMs through subsequent sequencing. Out of the 15.8K GEMs, GMM-Demux identifies and removes 2.8K multiplets, reducing the multiplet rate from 23.9% to 6.45%. After removing all the phony-type GEM clusters, GMM-Demux further reduces the multiplet rate to 3.29%.

## 2 Methods

GMM-Demux is built around four goals: 1) separate MSMs from SSDs in a sample barcoding dataset; 2) estimate singlet and SSM rates of a sample barcoding dataset; 3) plan future sample barcoding experiments: estimate the anticipated MSM, SSM and singlet rates of a planned future experiment; and 4) determine whether a homogeneous GEM cluster is a pure-type GEM cluster. GMM-Demux has two separate components: 1) a Gaussian-mixture-model-based MSM classifier and 2) a model based SSM rate estimator. The MSM classifier classifies GEMs into MSMs and SSDs using Gaussian-mixture models and computes the likelihood of each classification. The SSM rate estimator estimates the SSM and the singlet rate of the dataset. The SSM rate estimator models the GEM formation process as an augmented binomial process. It infers the latent parameters of the model, such as the number of cells of each sample and the number of cell-assay droplets formed during sequencing, from observed variables, including the number of cell-enclosing GEMs of each sample and the number of MSMs of each sample pair. Finally, the SSM rate estimator computes the estimated singlet and SSM rates of each sample with the inferred latent parameters. With the GEM formation model, GMM-Demux determines whether a proposed homogeneous GEM cluster is a pure-type GEM cluster, a phony-type GEM cluster or a mixture cluster.

Based on the GEM formation model, we build an online sample barcoding experiment planner that estimates the multiplet rates of future sample barcoding experiments. Researchers can use the experiment planner to anticipate the outcome of a sample barcoding experiment without actually conducting the experiment. The online experiment planner takes the number of cells planned for sequencing as well as the number of samples planned for sample barcoding as inputs and outputs the estimated MSM, SSM and singlet rates of the anticipated outcome.

### 2.1 Multi-Sample Multiplet (MSM) Classifier

The MSM classifier pre-processes the HTO matrix with centered-log-ratio (CLR) normalization[37, 36]. CLR normalizes the HTO UMI counts of each GEM column-wise (sample-wise) as follows:

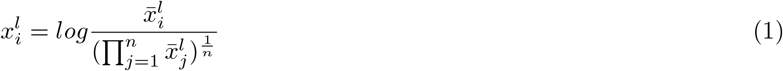

Here 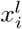 denotes the CLR-normalized HTO UMI count of the *l*th sample in the *i*th GEM (the *i*th row and the *l*th column of the HTO matrix); 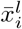 denotes the original HTO UMI count of the *l*th sample in the *i*th GEM and *n* denotes the total number of GEMs.

The distributions of the CLR-transformed HTO UMI counts of a 4-sample cell hashing experiment are illustrated in Figure 4. From this figure, we observe that for each sample, the CLR-transformed HTO UMI counts follow a bimodal distribution which resembles a mixture of two Gaussian distributions. GMM-Demux models the HTO UMI count distribution with an aggregated two-Gaussian-distribution mixed model. We color the two distributions as red and green respectively in Figure 4. For a specific sample 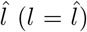, the Gaussian distribution with the smaller mean, 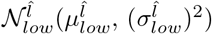 (in red), accounts for GEMs that do not contain cells from 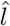 (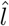-cell-free GEMs). The other distribution, 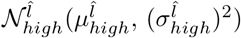 (in green), on the contrary, models GEMs that contain cells from 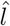 (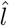-cell-enclosing GEMs). It is worth noting that GEMs from 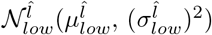 still have positive HTO counts. In cell hashing, when cell assays of all samples are pooled together, free-floating HTO antibodies that have not yet bound to any cell still exist in the solution, as shown in Figure 1. These residual free-floating HTO antibodies bind randomly to all cells from all samples (the restaining step in Figure 1). However, as cell assays are pooled together, antibodies are diluted, hence 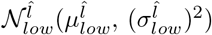 has a lower mean 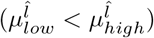. GEMs from 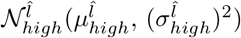, on the other hand, bind with HTO antibodies prior to pooling of samples. Before pooling, HTO antibodies have much higher concentrations. As a result, 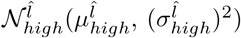 has a higher mean.

**Figure 4:**
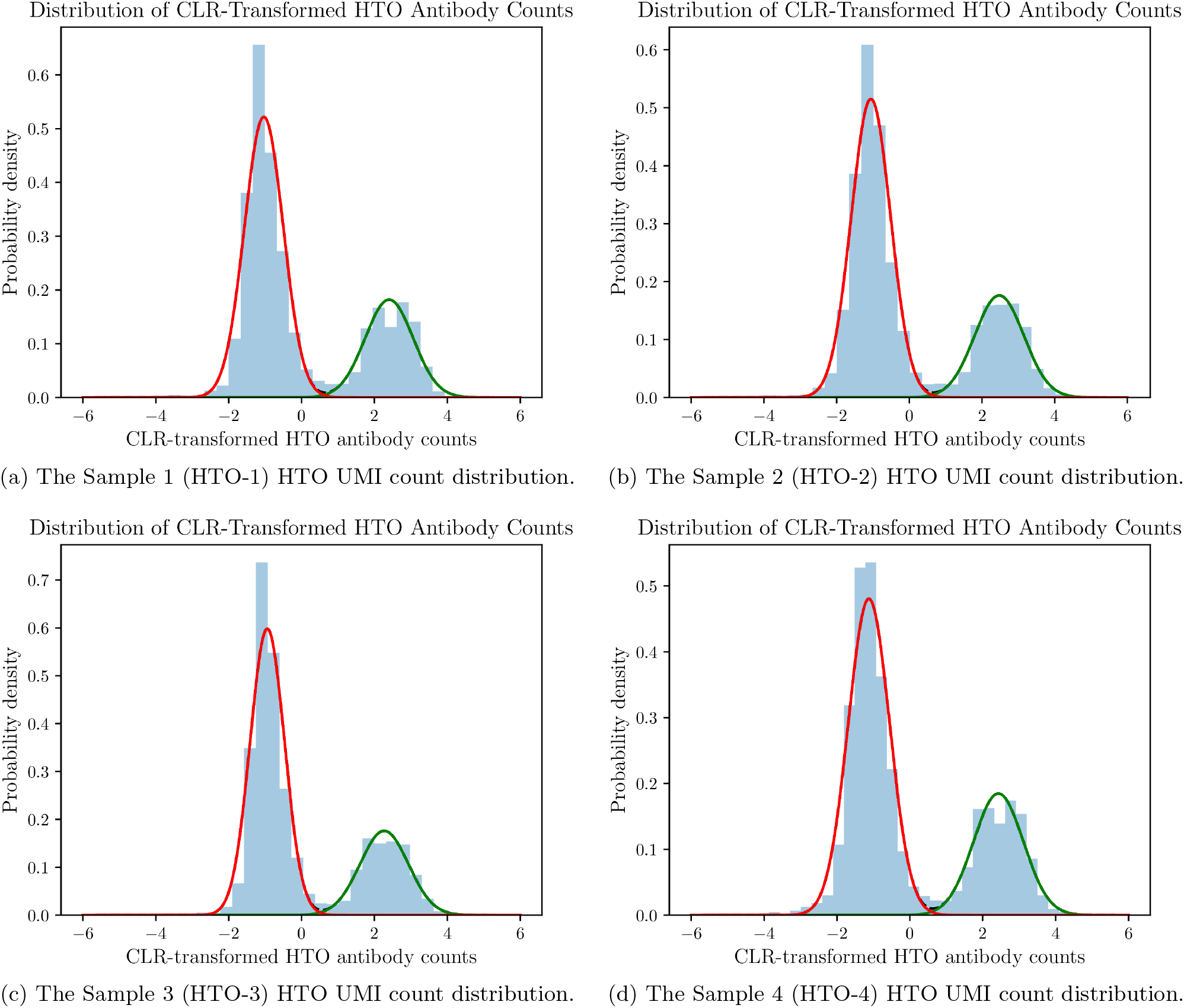
Distributions of CLR-transformed HTO UMI counts from a 4-sample cell hashing experiment. Each distribution is decomposed into two Gaussian distributions, 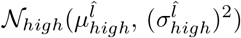 and 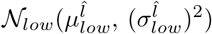. 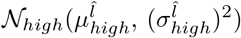 represents the HTO UMI count distribution of cell-enclosing GEMs. 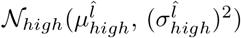 represents the HTO UMI count distribution of cell-free GEMs.

For each sample, GMM-Demux uses its Gaussian-mixture model to find GEMs that contain cells from the sample. Given a GEM, *i*, and a sample 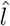, GMM-Demux tests whether 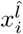 originates from the 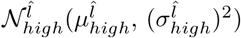 distribution of 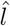: if 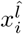 originates from 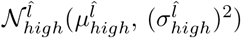, then *i* must contain cells from 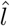; otherwise, 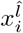 must belong to 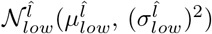, which means GEM *i* does not contain cells from 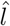.

Let 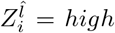 denote the event that 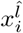 originates from 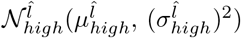 and 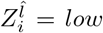 denote the event that 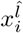 originates from 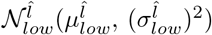. Let 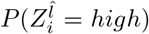 and 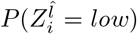 denote the prior probability of GEM *i* originating from 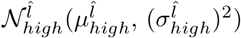 and 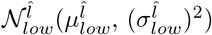 respectively. Then the probability of observing HTO count value 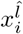 in GEM *i* equals to

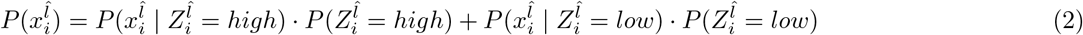

where 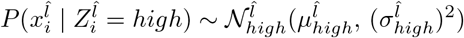 and 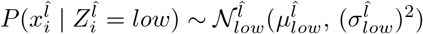.

GMM-Demux computes the mean and the standard deviation of 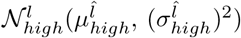 and 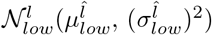; as well as the prior probabilities 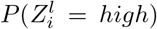 and 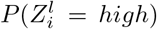 of each sample *l* using the Expectation Maximization (EM) Technique[34].

With all Gaussian-mixture models computed across all samples, for each GEM *i*, GMM-Demux computes the posterior probability of GEM *i* containing cells from sample 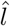. Let 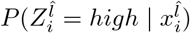 denote the posterior probability of 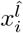 originating from 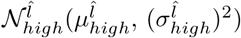, and 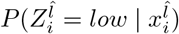 denote the probability of 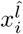 originating from 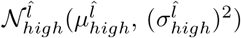. Both posterior probabilities 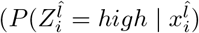 and 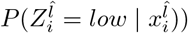 are computed using Bayes’ rule:

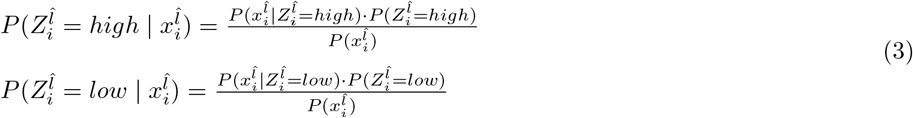

The probability 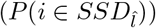 of *i* being a single-sample droplet (SSD) of sample 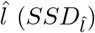 can be computed as:

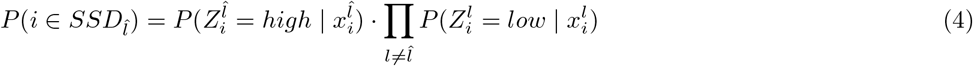

While the probability of *i* being a multi-sample multiplet (MSM) can be computed as:

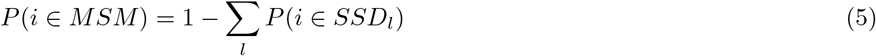

GMM-Demux classifies GEMs by ranking above probabilities: a GEM *i* is classified as a SSD of 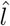 if 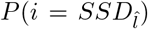 is the largest among all; or as a MSM if *P*(*i* = *MSM*) is the largest among all.

In fact, GMM-Demux is able to compute the probability of a GEM containing cells of any specific multi-sample configuration. Assume *U* is a set of samples (e.g., sample *l*_1_ and sample *l*_4_). The probability of GEM *i* containing cells from *U*, *MSM_U_* can be computed by:

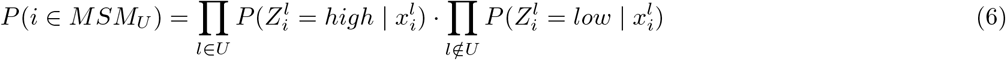

This allows GMM-Demux to not only identify and count SSDs, but also identify and count MSMs of specific sample combinations in a sample barcoding dataset. Counting MSMs of specific sample combinations is key to verifying the correctness of the SSM rate estimator, as we will show in later sections.

GMM-Demux lets the user specify a confidence cutoff c to filter out uncertain classifications. Sometimes GEMs have HTO UMI counts that reside in the junction area between 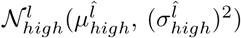 and 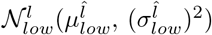 on a HTO sample dimension. Such GEMs produce ambiguous classification results: they have similar likelihoods between multiple classifications, which typically are all below 0.5. Uncertain GEMs are pruned by the confidence cutoff *c*: GEMs whose maximum probabilities across all classifications are less than *c* are deemed *uncertain GEMs* and are removed from the population. By tweaking *c*, GMM-Demux allows users to adjust the level of rigorousness in identifying SSDs and MSMs.

### 2.2 Same-Sample-Multiplet (SSM) Rate Estimator

As previously discussed, sample barcoding cannot distinguish SSMs from singlets. While GMM-Demux does not seek to identify SSMs in SSDs, it estimates the percentage of SSMs and singlets in each sample using the SSM rate estimator. Estimating the SSM rate in a dataset is critical for quality control. SSM rate represents the noise level of a sample. Samples with high SSM rates have low quality and should be removed.

GMM-Demux estimates the percentage of SSMs among all GEMs using a probabilistic model that models the entire GEM formation process in sample barcoding. The GEM formation process occurs after pooling of samples and governs the subsequent random distribution of cells into GEMs. GMM-Demux models the GEM formation process as an augmented binomial process: It assumes that after pooling of samples, the entire cell assay is divided into a finite number of droplets, called cell-assay droplets. Each cell is randomly and independently partitioned into a cell-assay droplet. During the single cell barcoding process, a fraction of all cell-assay droplets are combined with gel beads and form GEMs. The rest of the cell-assay droplets do not form GEMs and will not be sequenced. We use the term *droplet capture rate* to denote the probability that a cell-assay drop is combined with a gel bead. GEMs, which contain both cell-enclosing cell-assay droplets and gel beads, are recovered after sequencing and are summarized in a HTO matrix. A detailed illustration of the GEM formation model is provided in the Supplementary Materials, in Section S1.

The rates of multiplets, including both SSM rates and MSM rates, are modeled as the probability of having multiple cells (from the same or different samples) partitioned into the same cell-assay droplet. A major challenge for this method is that key parameters, namely the number of cells in each sample, the droplet capture rate, and the total number of cell-assay droplets, are not directly observable. Instead, from the MSM classifier, we observe the number of sample-specific GEMs as well as the number of MSMs of any sample pair. Combined with the prior knowledge of the estimated total number of cells loaded for sample barcoding, the SSM rate estimator derives the latent parameters of the model and uses the complete model to estimate the multiplet rates of the dataset.

#### 2.2.1 Modeling Multiplets

The SSM rate estimator models the GEM formation process as follows: Assume there are a total of *X* cell-assay droplets. Also assume there are *y_l_* cells in a sample, *l*, with *Y* denoting the overall population of all cells, or *Y* = Σ_*l*_ *yl*. The model assumes that each cell is independently and randomly partitioned into a cell-assay droplet. Consequently, a cell has a probability of 1/*X* to reside within a specific cell-assay droplet. Assuming that no bias exists among cells from different samples, then the probability of a cell-assay droplet, *i*, being a singlet, given that *i* is not empty, can be calculated as

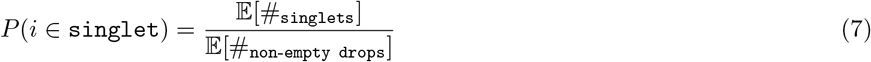

where 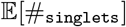 is the expected number of singlets and 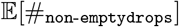 is the expected number of non-empty cell-assay droplets. For simplicity, in the rest of this paper, we refer to cell-assay droplets simply as droplets.

Since cells are randomly partitioned into droplets, 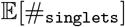 can be computed from a binomial model. Specifically we have 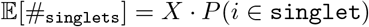, where *P*(*i* ∈ `singlet`) denotes the probability of having one and only one cell, out of a total of *Y* cells, residing in *i*. All other cells are partitioned into other droplets. Mathematically, we have:

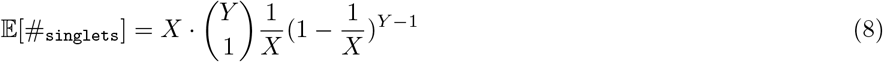

Similarly, the expected number of non-empty droplets can be computed as 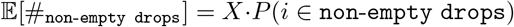. *P*(*i* ∈ `non-empty drops`) is the probability of droplet *i* being non-empty and it equals to 1 − *P*(*i* ∈ `empty drops`). According to binomial distribution, *P*(*i* ∈ `empty drops`) equals to the probability of all cells residing in droplets other than *i*. Overall, we have:

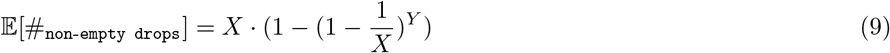

Equally, the probability of *i* being a MSM given *i* is not empty, *P*(*i* ∈ `MSM`), can be computed as

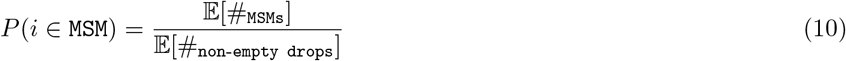

with 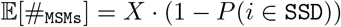, where *P*(*i* ∈ `SSD`) is the probability of *i* being a SSD.

When more than one sample is labeled in sample barcoding, we have *P*(*i* ∈ `SSD`) = Σ_*l*_ *P*(*i* ∈ `SSD`_*h*_), where *P*(*i* ∈ `SSD`_*l*_) is the probability of *i* being a SSD of sample *l*. Let set *D_l_* represent all and only *l*-cell-enclosing droplets and set 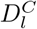 to represent all and only *l*-cell-free droplets. The probability of droplet *i* being a SSD of sample 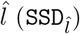 equals the probability of *i* being a cell-enclosing droplet in 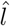 and a cell-free droplet in all other samples. Based on binomial distribution, the probability of *i* belonging to 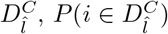, equals the probability of all cells of 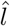 residing in droplets other than *i*, which is 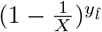. As 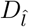 and 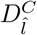 complements each other, we have 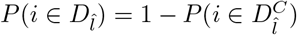. We expand 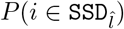 into the following:

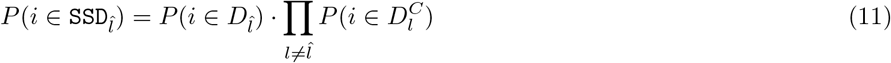

where 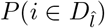 and 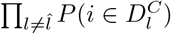 can be computed as:

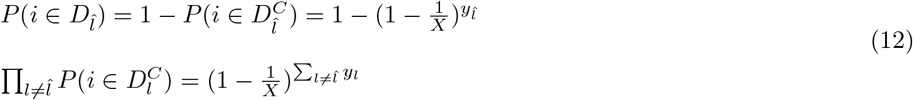

Finally, the probability of *i* being a SSM is simply the probability of *i* being neither a MSM nor a singlet. Mathematically, we have:

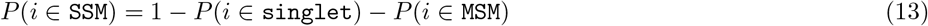

Alternatively, we can compute *P*(*i* ∈ `SSM`) as *P*(*i* ∈ `SSM`) = Σ_*l*_ *P*(*i* ∈ `SSM`_*l*_), with *P*(*i* ∈ `SSM`_*l*_) denoting the probability of *i* being a SSM of sample *l*. Because a SSD of *l* must either be a SSM of *l* or a singlet of *l*, therefore event {*i* ∈ `SSM`_*l*_ | *i* ∈ `SSD`_*l*_} and event {*i* ∈ `singlet`_*l*_ | *i* ∈ `SSD`_*l*_)} must be collectively exhaustive events. Together, *P*(*i* ∈ `SSM`_*l*_) can be computed as:

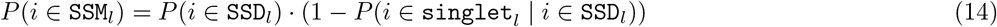

Since all singlets of *l* are SSDs of *l*, we have:

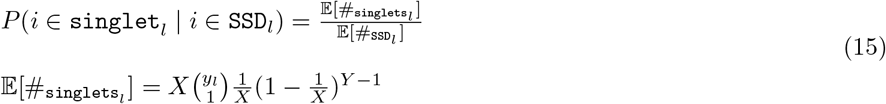

The two methods (Equation (13) and (14)) of calculating *P*(*i* ∈ `SSM`) are equivalent (details are omitted to conserve space).

Overall, given *X* and *y_l_* for every sample *l* of a sample barcoding dataset, the SSM rate estimator estimates the singlet rate (*P*(*i* ∈ `singlet`), Equation (7)), the MSM rate (*P*(*i* ∈ `MSM`), Equation (10)) and the SSM rate (*P*(*i* ∈ `SSM`), Equation (13) and Equation (14)) of the dataset. Unlike the SSM rate, which can only be inferred indirectly through the GEM formation model; the MSM rate can be obtained both analytically through the GEM formation model and numerically by interpreting the MSM classification result. As a result, we can validate the correctness of the GEM formation model by comparing the MSM rates obtained through both methods. In the Result section, we show that both methods provide consistent MSM rates.

We perform simulations to verify the correctness of the above equations. The simulation results are included in the Supplementary Materials, in Section S2. Specifically, we repeatedly simulate the GEM formation process. We show that the singlet, SSM and MSM rates measured from simulations asymptotically match the values analytically computed with above equations.

#### 2.2.2 Estimating Model Parameters

GMM-Demux relies on *X* and *y_l_* of every sample *l* to compute the SSM rates. However, neither *X* nor *y_l_* is directly observable in a sample barcoding dataset. Instead, from the classification result, GMM-Demux observes *z_l_*, the number of GEMs in *D_l_*.

Let *r_cap_* denote the droplet capture rate. From *z_l_* and a user-provided estimation of the total cell count, *Y*, GMM-Demux computes both *X, r_cap_* and *y_l_*. For a HTO sample *l*, based on our multiplet model, we have 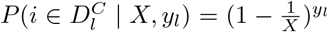 (*X* and *y_l_* serve as parameters) and 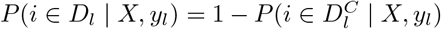. Let random variable *Z_l_* denote the number of GEMs that enclose cells from *l* and let *P*(*Z_l_* = *z_l_* | *X, r_cap_, y_l_*) denote the probability of observing *z_l_ l*-cell-enclosing GEMs under the parameter set [*X, r_cap_, y_l_*]. According to the GEM formation model, which models partitioning of cells into droplets with a binomial distribution, we have:

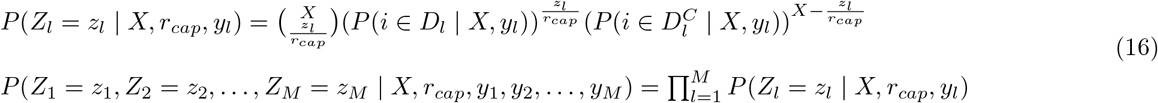

We derive the model parameters by computing

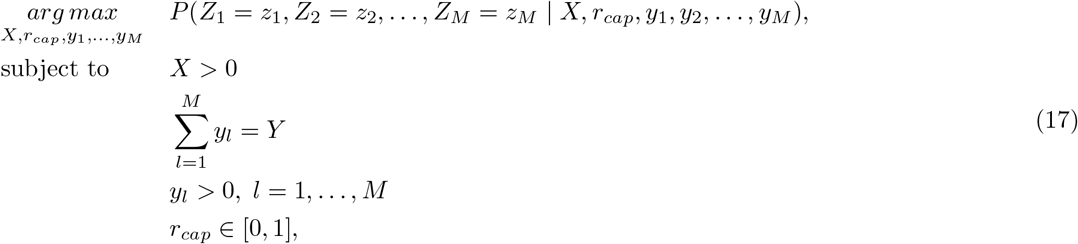

where *Y* is the user-provided total number of cells loaded for library prep, which can be obtained from the hemocytometer.

### 2.3 Online Cell Hashing Experiment Planner

The online sample barcoding experiment planner estimates the singlet, SSM and MSM rates of a planned sample barcoding experiment via the GEM formation model. Specifically, it takes the estimated number of cells (*Y*), the planned number of samples for sample barcoding (*M*), the estimated number of droplets (*X*), and the droplet capture rate (*r_cap_*) in library prep as inputs, and it computes the estimated multiplet rates. The online experiment planner assumes cells are evenly distributed among *M* samples.

The online experiment planner also estimates the relative single-sample-multiplet (RSSM) rate, defined as the estimated number of SSMs among SSDs. Mathematically, RSSM rate is defined as

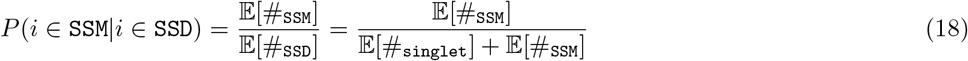

The RSSM rate marks the overall quality of a sample barcoding dataset. It represents the percentage of irremovable multiplets among SSDs, after removing all MSMs in the dataset. If the RSSM rate of the estimated outcome is too high, then the planned experiment should be aborted, as the anticipated outcome will be too noisy for downstream analysis. While dividing the cell assay into more samples drives down the RSSM rate, as it reduces 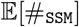, it increases both the cost and the complexity of the experiment. With the multiplet rate estimator, researchers can determine the minimum number of HTO samples to use in a sample barcoding experiment, to save cost while meeting the RSSM rate target.

The online experiment planner computes the multiplet rates as follows:

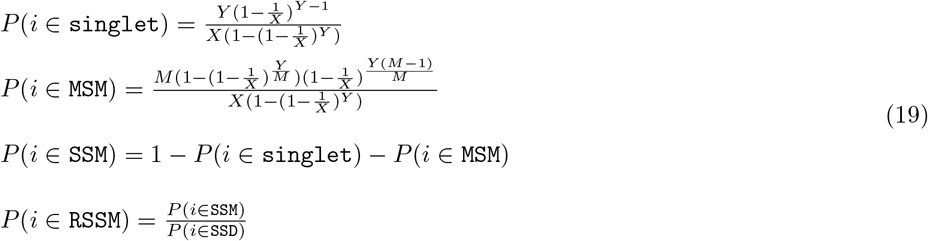

The above equations show that the number of samples, *M*, does not affect the singlet rate. The singlet rate is solely determined by *X* and *Y*. However, a greater *M* reduces the SSM rate and increases the MSM rate. Therefore, we conclude that dividing a cell assay into more samples by sample barcoding transforms more SSMs into MSMs. Transforming SSMs into MSMs improves the quality of the dataset. With fewer SSMs, the RSSM rate of the dataset decreases. In comparison, having more MSMs does not affect the quality of the dataset, as MSMs are removed by GMM-Demux.

Given *r_cap_*, the online experiment planner also computes the estimated number of cell-enclosing GEMs in the final output, as well as the estimated number of SSDs after removing MSMs. The number of cell-enclosing GEMs, #_non-empty GEM_ = #_non-empty drops_ · *r_cap_*(#_non-empty drops_ is computed in Equation (9)). The number of SSDs is computed as #_SSD_ = #_non-empty GEM_ · (1 − *P*(*i* ∈ `MSM`)).

Among all four inputs, *Y* and *M* are user-controlled while *X* and *r_cap_* are largely dictated by the library prep equipment. However, based on our observations, we found that *X* mostly varies between 65K-80*K*. To account for the wide ranges of variability of the inputs, the online experiment planner uses sliders for selecting *X, Y, M* and *r_cap_*, which have ranges of 60*K*-100_K_, 1*K*-80*K*, 1-20 and 0 − 1 respectively. The online experiment planner supports dynamic updates. It computes the estimated multiplet rates in real time as the user updates input parameters. In practice, we recommend that users profile their library prep equipment once for the total number of droplets (*X*) in a sequencing run, by performing a small-scale sample barcoding experiment and use the profiled *X* (included in the GMM-Demux output) in planning future experiments.

### 2.4 Pure-Type GEM Verification

In novel cell type identification, a cell type classifier is used to group GEMs into clusters. Each cluster is assumed to represent a unique cell type. Clusters whose average expression profiles do not match any known cell types are identified as novel cell types[40].

After clustering, phony-type GEMs are grouped into distinct clusters. Phony-type GEM clusters may be incorrectly identified as novel cell types, as their expression profiles do not match known cell types, generating false discoveries. GMM-Demux rectifies true novel cell types by validating if the alleged novel-cell-type GEM cluster contains mainly pure-type GEMs. Based on the GEM composition in the cluster, GMM-Demux classifies GEM clusters into three categories: pure-type GEM clusters, phony-type GEM clusters and mixture clusters. Phony-type GEM clusters contain mostly phony-type GEMs.Pure-type GEM clusters contain mostly pure-type GEMs.Mixture clusters contain large quantities of both pure-type and phony-type GEMs.

Let *G* represents a GEM cluster. GMM-Demux classifies *G* by examining the MSM ratio of G. For simplicity, we assume cells are equally randomly divided into the *M* sample barcoding samples. If *G* is a phony-type GEM cluster, the MSM ratio of *G* must be very high. Elaborated in the Supplementary Material section 3, the expected MSM ratio of a phony-type cluster approaches and exceeds 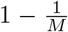. Otherwise, if *G* is a pure-type GEM cluster, its MSM ratio should not be greater than the MSM ratio of the entire sample barcoding dataset, which is much smaller than 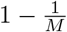. The MSM ratio reflects the GEM composition of *G*: in a phony-type GEM cluster, all GEMs are multiplets, hence the MSM ratio of *G*, *r*_MSM_G__, equals to 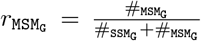, where #_MSM_G__ and #_SSM_G__ denote the number of MSMs and SSMs in *G*, respectively; in a pure-type GEM cluster, however, we have 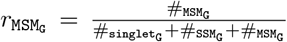 instead, where #_singlet_G__ denotes the number of singlets in *G*. By comparing the two ratios, we observe that pure-type GEM clusters include singlet counts in the denominator, whereas phony-type GEM clusters do not. As a result, the MSM ratio is much higher in phony-type GEM clusters than in pure-type GEM clusters. Complex situations where cells are not evenly distributed among sample barcoding samples are discussed in the Supplementary Materials, section 3.

GMM-Demux uses hypothesis testing to measure the confidence of each classification. GMM-Demux prepares two hypotheses, the *phony-type hypothesis* and the *pure-type hypothesis*, which assume *G* being a pure-type or a phony-type GEM cluster, respectively. GMM-Demux tests both hypotheses with the binomial test and computes a p-value for each hypothesis. Based on the hypothesis testing results, GMM-Demux classifies *G* as a pure-type GEM cluster, a phony-type GEM cluster or a mixture cluster. Details of the hypothesis tests are provided in the Supplementary Materials, section 3.

Based on the classification result of *G*, GMM-Demux recommends different actions. Being classified as a phony-type GEM cluster suggests that the proportion of pure-type GEMs in *G*, if there exists any, is extremely small and most GEMs in *G* are phony-type GEMs. GMM-Demux recommends excluding *G* from further analysis.Being classified as a mixture cluster suggests that *G* mixes pure-type GEMs and phony-type GEMs together and has non-trivial numbers of GEMs in both categories.This is often a result of poor clustering quality where *G* becomes a super-cluster over several pure-type and phony-type GEM clusters. GMM-Demux recommends refinement over the clustering method and subdividing *G* into pure-type GEM and phony-type GEM sub-clusters. Finally, being classified as a pure-type GEM clusters suggests that it is plausible that *G* defines a real cell type. Further analysis over *G* is recommended.

### 2.5 Compatibility

The GMM-Demux classifier is compatible with CellRanger-3.0 from 10X Genomics. It takes the sample barcoding data, in the market matrix (mtx) format, together with the estimated number of cells (*Y*), as inputs and it outputs a double column table as the classification result. The row indices of the output table are GEM barcodes. The two columns are the classification of each GEM and the confidence score of each classification, respectively. With *M* samples, GMM-Demux classifies GEMs into a maximum of 2^*M*^ + 1 classes. Besides the uncertain class, the negative class and *M* SSD classes, there are 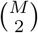 bi-sample classes, 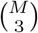 tri-sample classes,… and 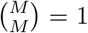 *M*-sample class. Additionally, GMM-Demux produces a SSM-rate summary file, which includes the SSM rate and the RSSM rate of each sample, and a summary file that includes the multiplet rates of the entire dataset. The summary file also includes the estimated number of cell-assay droplets (*X*) and the estimated droplet capture rate (*r_cap_*) of the library prep equipment. Example outputs are provided in the Supplementary Materials, in Section S4.

## 3 Related Works

The heuristic classifier currently is the only analytical tool for processing sample barcoding data. It relies on the K-medoid clustering algorithm[10], a probabilistic method[31], to classify MSMs. Assuming there are a total of *M* samples, for each sample, it clusters all GEMs into *M* groups using the K-medoid clustering algorithm. Then it removes the group with the highest mean; combines the remaining groups; fits the combined data with a negative binomial distribution; excludes the top 5% values as outliers; computes the *q* = *t_l_* quantile (*t_l_* is set to 99% by default) of the fitted distribution; and finally tag GEMs whose HTO UMI values that are greater than *q* as sample-specific GEMs. If a GEM is tagged as sample-specific by multiple samples, then the heuristic classifier brand it as a MSM.

While the heuristic classifier is sufficient to demonstrate the benefit of sample barcoding, it is heuristic-based and is unstable. It includes a number of arbitrary parameters. It does not explain why it fits the data with a negative binomial distribution as opposed to other distributions; nor does it explain why it removes the top 5% values as outliers or sets *t_l_* = 99% as the default value. As we will see in the Result section, by setting *t_l_* differently, it generates conflicting results and it is not evident which *t_l_* provides the best result. Furthermore, because it relies on the K-medoid clustering algorithm, which generates inconsistent results over repetitive runs, the heuristic classifier also generates inconsistent classification results over repetitive executions.

Finally, the heuristic classifier does not model the GEM formation process. As a result, it is incapable of estimating the percentage of SSMs among SSDs. Consequently, it cannot grade the quality of a sample barcoding dataset, nor can it guide future sample barcoding experiment designs.

Prior to sample barcoding, multiplets can be identified experimentally by mixing samples of different donors. The most reliable method of finding multiplets involves mixing cells of different species[45, 18, 13, 6]. Multiplets are identified as GEMs whose reads are confidently mapped across multiple species. However, this method does not work when mixing samples of the same species. Instead, when working with samples of the same species, as long as the donors show sufficient amount of genetic variations, then multiplets can be identified as GEMs who contain distinct genetic signatures from multiple donors[12]. Unfortunately, neither method works when samples come from a single donor, which limits their applicability in scaling up single cell experiments. Sample barcoding, on the other hand, is capable of identifying multiplets even when samples are drawn from the same donor.

Besides the aforementioned methods, it is also plausible to identify some doublets through examining single cell expression profiles alone. When working with multiple purified cell types, under the assumption that cells of the same type have highly similar expression profiles while cells of different cell types have drastically different expression profiles, multiplets are identified as small GEM groups whose expression profiles share similarities to multiple distinct large GEM groups or to multiple expression profiles of known distinct cell types[45, 36]. This idea can be further expanded into artificially create synthetic doublets from a single cell dataset and detect doublets by selecting GEMs whose expression profiles resemble synthetic doublets[44, 22]. While the idea is promising, it has a few critical limitations. The entire idea was built on the presumption that cells of different types have distinctive expression profiles. As a result, not only can it not find multiplets formed by cells of the same type, it also cannot detect multiplets comprised of cells from similar lineages, such as CD4^+^ T cells and CD8^+^ T cells. Furthermore, it cannot reliably distinguish multiplets from transitional cells, as transitional cells also have expression profiles that are similar to multiple distinct mature cell types[35]. In this paper, we focus on analytically detecting multiplets through sample barcoding.

There are only a few prior studies on modeling multiplet rates. Demuxlet[12], a genetic-variation based multiplet identifier, models the singlet rate as 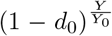, where *Y* is the planned number of cells and *d*_0_ is the observed doublet rate (through a mixed-species experiment) when loading Yocells in library prep. By default, Demuxlet assumes *d*_0_ = 0.01 with *Y*_0_ = 1*K*. Although not elaborated in the Demuxlet paper, we notice that the singlet rate equation in Demuxlet bears a striking resemblance to the singlet rate equation used by GMM-Demux. Specifically, within the range of *Y* ∈ [1*K*, 40_K_], 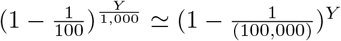. This is because the curve 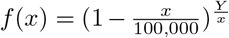 is almost flat within *x* ∈ [1,10, 000]. Hence, the singlet formula used by Demuxlet under *d*_0_ = 0.01 and *Y*_0_ = 1*K* can be approximately explained by GMM-Demux as randomly partitioning *Y* cells among a total of *X* = 100*K* cell-assay droplets. Despite apparent similarities between their formulas, GMM-Demux and Demuxlet employ different underlying statistical mechanics. Demuxlet uses a discriminative model, which uses regression to subjectively model the multiplet rate as a parametrized curve. GMM-Demux, on the other hand, uses a self-explanatory, generative model that directly simulates the GEM formation process. The generative model allows GMM-Demux to estimate the MSM rates of pure-type and phony-type GEM clusters in a sample barocding dataset, while the discriminative model of Demuxlet does not. The generative model also enables GMM-Demux to accurately simulate multiplets, including both pure-type and phony-type GEMs; singlets, SSMs and MSMs; whereas Demuxlet cannot.

Alternatively, other works model the number of cells in a GEM with Poisson distributions[7, 24, 3]. A major downside of this branch of methods is the difficulty in estimating the model parameters. A Poisson model uses the average number of cells in a GEM as its parameter. However, this number changes when the number of loaded cells changes. As a result, these models cannot be readily used for experiment planning. Interestingly, Poisson distribution is a special case of the binomial distribution, where the number of probabilistic experiments in the binomial process (*X*, in this case) approaches infinity[27]. Poisson distribution is often used as a numerically approximation of binomial distributions, especially when the number of droplets (X) is large and the average number of cells in a droplet is small. Poisson-distribution based multiplet rate estimators in fact supports the GEM formation model of GMM-Demux and can be considered as numerical approximations of GMM-Demux.

## 4 Datasets

### 4.1 Real Datasets

We benchmark GMM-Demux on three separate HTO datasets from three independent sources. In addition to a public dataset from Stoeckius *et al*.[37], we conducted two additional in-house cell hashing experiments independently in two separate labs. A summary of the three datasets is provided in Table 2.

**Table 2:** Summary of cell-hashing datasets.

| Name | Est. #Cells | #Cell-Assay Droplets | #Samples | Tissue | Source |
| --- | --- | --- | --- | --- | --- |
| PBMC-1 | 35,685 | 15,841 | 4 | PBMC | In house |
| CD4<sup>+</sup> Memory T | 25,000 | 9,715 | 5 | CD4<sup>+</sup> Memory T Cells | In house |
| PBMC-2 | 28,000 | 15,455 | 8 | PBMC | Stoeckius *et al.* [37] |

Cells in the PBMC-1 dataset are drawn from a healthy donor described in a previous study[38]. These cells are divided into four samples. Each sample is subjected to the Totalseq-A and cell hashing protocol[37], targeting a recovery of ~5,000 cells per sample. All HTO-tagged cells are pooled together and are prepared using the 10x Genomics platform with Gel Bead Kit V2. The prepared assay is subsequently sequenced on an Illumina Hiseq with a depth of 50K reads per cell. In addition to cell hashing, cells in this dataset are simultaneously measured for their surface marker abundance through CITE-seq[36]. Eight surface markers are measured for every cell: CD3, CD4, CD8, CD11, CD14, CD16, CD19 and CD56.

Human CD4^+^ Memory T cells were enriched from the peripheral blood of a healthy adult human volunteer using the MACSxpress®Whole Blood CD4 Memory T Cell Isolation Kit, human (MiltenyiBiotec). The cells were then incubated for 12 hours at 37°C, 5% CO_2_ and at a concentration of 1 ×10^−6^ cells/mL in serum-free, X-VIVO-20 medium (Lonza BioWhittaker) with T cell activation beads coated with anti-CD2/-CD3/-CD28 antibodies (Miltenyi Biotec) alone or in combination with four different sets of recombinant human inflammatory mediators (i.e., five different culture conditions). The cells were then harvested from the culture medium for cell hashing[37] and CITE-seq[36] single cell sequencing library preparation following the CITE-seq and hashing protocol available at https://cite-seq.com. The mRNA-, HTO-, and ADT-derived libraries were then pooled at approximately 85%, 5% and 10% proportions, respectively, and the pool of these sequencing libraries was sent for 150 bp paired end sequencing in two lanes of an Illumina HiSeq sequencer (MedGenome, Inc.).

All subjects were given informed consent and the study is approved by the University of Pittsburgh IRB.

### 4.2 Simulation Dataset

We also generated a simulated dataset by augmenting the PBMC-1 dataset. Specifically, we classify GEMs in the PBMC-1 dataset using both GMM-Demux and the heuristic classifier. Then we extract GEMs that are classified as SSDs by both classifiers. We assume these GEMs are SSDs in truth. We further assume there are a total of *X* = 80*K* cell-assay droplets with a capture rate *r_cap_* of 1. We randomly distribute the SSDs among the 80K droplets. If a droplet is assigned with a single SSD, then it inherits the HTO counts of that SSD. If a droplet is assigned with more than one SSDs, then the new HTO counts of the droplet is computed by adding the HTO counts of its assigned SSDs together. Let *j* denote a simulated multi-SSD droplet and 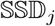 denote the set of SSDs assigned to *j*, we compute the new HTO counts of *j* as 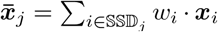, where *w_i_* is a random number generated from 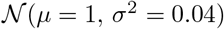 and ***x***_*i*_ is the HTO count vector of SSD *i*. Simulated multi-SSDs droplets that contain SSDs from multiple samples are marked as MSMs in ground truth. A perfect classifier should identify all MSMs without misclassifying SSDs into MSMs.

## 5 Results

### 5.1 Multi-Sample Multiplet Classification Results

For each cell-hashing dataset, we compare the classification results of three separate approaches: the GMM-Demux classifier, the heuristic classifier and a human-supervised classifier. For the human-supervised classifier, a trained laboratory technician classifies GEMs based on the CLR-transformed HTO matrix.

The classification results are visualized in 2D tSNE plots[16]. The tSNE plots are generated directly from the HTO matrix. Each dot in the tSNE plot represents a GEM. The color of the GEM specifies the class of the GEM. By the nature of tSNE plots, GEMs with similar HTO UMI count profiles are more likely to be drawn close to each other. Therefore, GEMs with the same classifications usually form natural clusters. Note that tSNE is probabilistic and non-deterministic: GEMs with similar HTO UMI count profiles are likely to be grouped together but there is no guarantee[42]. Sometimes, a small fraction of GEMs are incorrectly clustered with dissimilar neighbors, due to inaccuracies of the tSNE transformation. We use tSNE plots only for visualization and do not expect it to 100% reflect the truth.

#### 5.1.1 Classification Results on Real Datasets

The classification results of the real datasets are shown in Figure 5. For each dataset, we show three sets of figures: the GMM-Demux classification result in the top left panel; the heuristic classification result in the top middle panel; the human supervised classification result in the top right panel; and a set of HTO-UMI-count heat maps of individual samples in the bottom panel. In each heat map, GEMs with higher HTO UMI counts of the sample have darker colors. For simplicity, we lump all MSMs together as a single class—the MSM class, while maintaining SSDs of different samples as separate classes. If needed, GMM-Demux is able to subdivide MSMs into sub-classes where each sample combination is given a distinct class. Distinct MSM classification results are provided in the Supplementary Materials in Figure S2.

**Figure 5:**
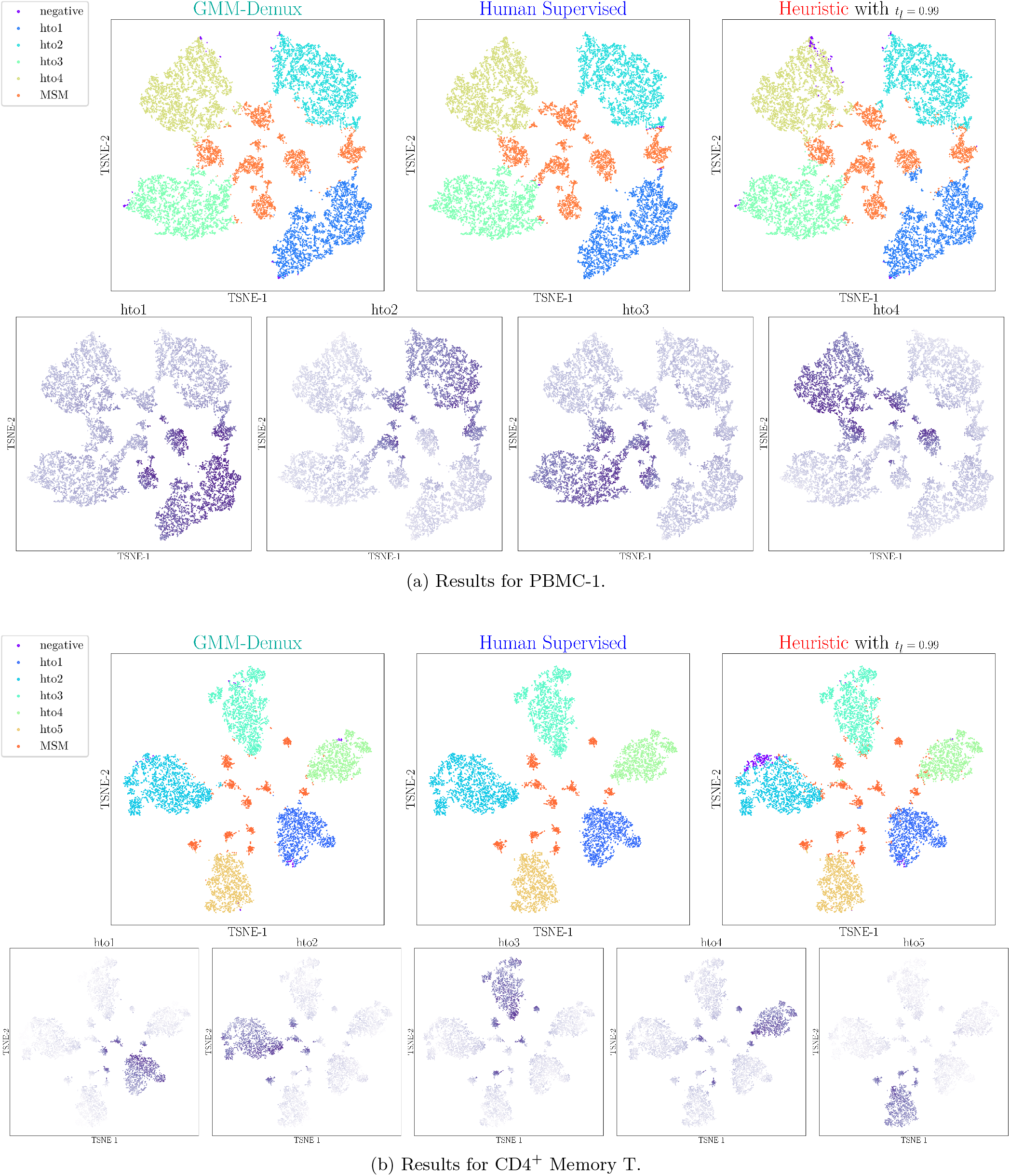
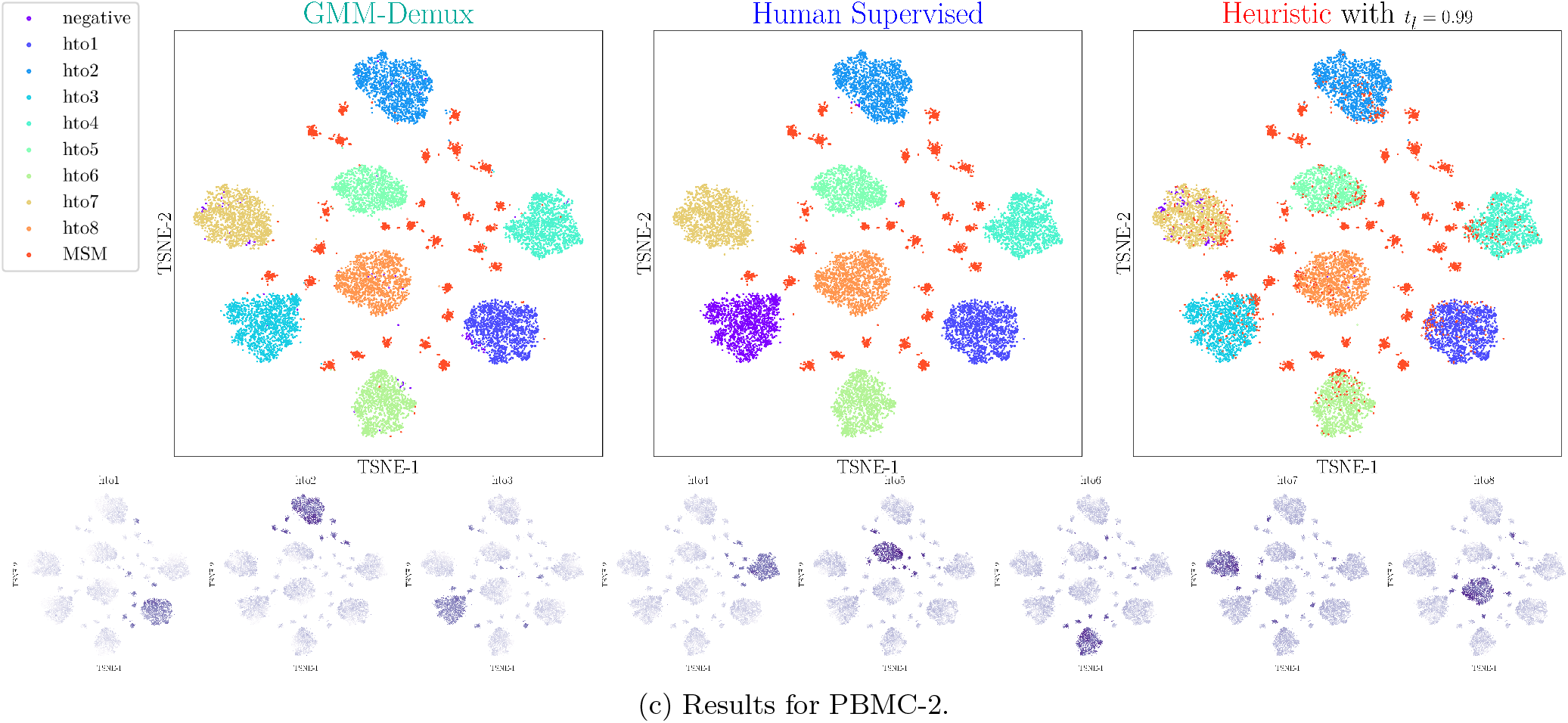
Classification results for each dataset. Each dot represents a cell. For each dataset, the upper panels present classification results produced by different classifiers. The lower panel stores the heat maps of HTO UMI counts of individual samples.

Figure 5 shows that the classification results from all three classifiers are mostly consistent. We compare the classification results against the HTO-UMI-count heat maps: a correct SSD classification should have dark color in a single heat map and light colors in the rest heat maps; a correct MSM classification should have dark colors in more than one heat maps. Among all three classifiers, GMM-Demux adheres to the HTO heat maps the most. The human-supervised classifier is the most prone to classification errors that stem from inaccuracies in tSNE transformations. During manual classification, GEMs that are close-by in tSNE plots are often assumed to be similar, which is an unreliable assumption, due to limited accuracy of the tSNE transformation.

Even though the heuristic classifier generates classification results similar to those produced with the GMM-Demux classifier, it is heuristic-based and unstable. Figure 6 illustrates the heuristic and unstable nature of the heuristic classifier. Results in this figure are generated from the PBMC-1 dataset. Since the heuristic classifier relies on the HTO UMI count threshold for classification, which is indirectly controlled by *t_l_*, it generates different classification results with different *t_l_* values, as shown in Figure 6a-6d. From above figures, we observe that while a smaller *t_l_* produces fewer negative classifications, it generates more MSM classifications. This is expected as a smaller *t_l_* reduces the HTO UMI count threshold, which in turn increases the number of cell-enclosing GEMs in each sample. Without ground truth, however, it is not obvious which *t_l_* provides the most accurate classification result. Such high variations in classification results, as well as the heavy reliance on heuristic parameters, reduce the reliability of the heuristic classifier. In practice, it is difficult to select the appropriate t_l_ for the best accuracy.

**Figure 6:**
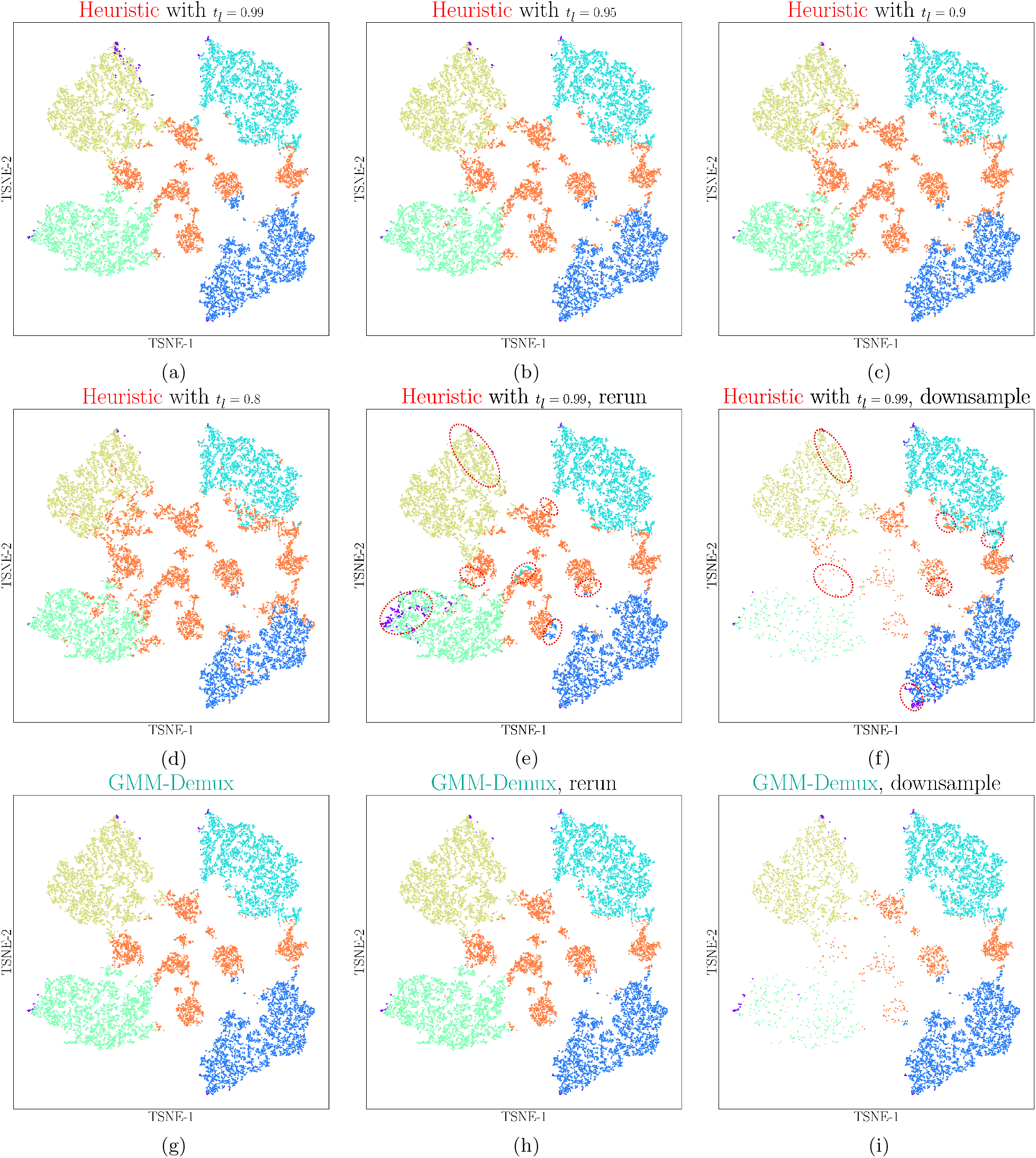
Stability test results. The heuristic classifier produces different classification results with regard to varying heuristic parameters, *t_l_*. It also generates inconsistent classification results during repetitive executions and is susceptible to data augmentation (sub-sampling). Classification differences of the heuristic classifier are highlighted in red-dotted circles. GMM-Demux, on the contrary, generates consistent classification results.

On top of its heuristic nature, because it uses the non-deterministic K-medoid clustering algorithm, the heuristic classifier generates different results between two runs even with the same heuristic parameter. This is visualized by comparing Figure 6a against Figure 6e. Both figures are generated under *t_l_* = 0.99. Differences between them (highlighted in red-dotted circles) stem solely from the non-determinism of the K-medoid algorithm.

Finally, the heuristic classifier is highly sensitive to changes in the dataset. In Figure 6f, we randomly sub-sample GEMs from sample 3 and 4 (by 10% and 50% respectively). When compared against Figure 6a, we observe substantial changes in the classification result, highlighted in red-dotted circles. This is because as the sample composition changes, the HTO count threshold of each sample also changes, even without updating *t_l_*. As a result, previously-classified MSMs now become SSDs and vice versa.

The GMM-Demux classifier, on the other hand, is model-based, stable and far more deterministic. The GMM-Demux classifier does not require heuristic parameters for MSM classification and generates consistent classification results across repetitive runs. Despite of uncertainties introduced by the EM algorithm, because GMM-Demux is model-based and the HTO UMI count distributions possess obvious features of a 2-component Gaussian mixture, the EM algorithm always converges. Hence, GMM-Demux generates consistent results. Figure 6g and Figure 6h show the classification results of two repetitive runs of GMM-Demux. There exist little differences between the two figures. Similarly, the GMM-Demux classifier is much less susceptible to sub-sampling, as shown in Figure 6i, where we sub-sampled GEMs from sample 3 and 4, as we did in Figure 6f. By comparing Figure 6i against Figure 6g, we observe minimal changes in GEM classifications.

Not all GEMs can be confidently classified by GMM-Demux. Some GEMs have low HTO UMI counts across all samples; while other GEMs have similar probabilities between multiple classes (such as between a *l*_1_ BSD and a *l*_1_ ∩ *l*_2_ MSM). Neither type of GEMs can be well classified: the former are classified as *negative* GEMs, which should be experimental errors; while the latter are classified as *unclear* GEMs, which are too ambiguous to be included in the final result. GMM-Demux lets the user specify the confidence threshold, *c*, such that the user can customize the removal of unclear GEMs: a low confidence threshold salvages more unclear GEMs in the final result at the expense of decreased classification quality. Across all three cell hashing datasets, over 99% of GEMs have confidence scores above 0.8. Therefore, we set the default confidence threshold of GMM-Demux at 0.8 (*c* = 0.8). Detailed distributions of confidence scores are provided in the Supplementary Materials, in Figure S3.

#### 5.1.2 Classification Results on the Simulation Dataset

We compare the accuracy of GMM-Demux against the heuristic classifier by applying both methods to the simulation dataset and compare their classification results against the ground truth. Figure 7 shows the classification results on the simulated dataset. Compared to the heuristic classifier, which mis-classifies 4.571% GEMs, GMM-Demux mis-classifies only 0.066% GEMs, resulting in a 69× reduction in classification error rate.

**Figure 7:**
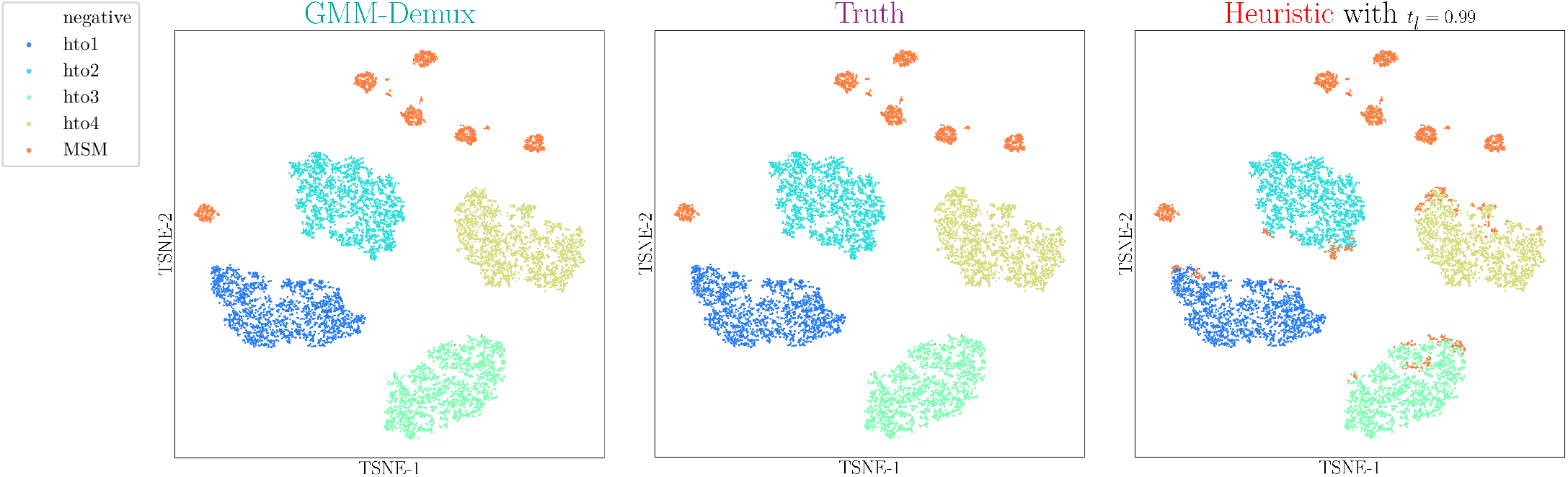
Classification results on the simulated dataset. Compared to truth, the heursitic classifier mis-classifies 4.571% GEMs while GMM-Demux mis-classifies 0.066% GEMs.

### 5.2 Same-Sample Multiplet Rate Estimation Results

We prove the correctness of the SSM estimator indirectly by validating the GEM-formation model. Even though the SSM rate truth is not directly observable, if the underlying probabilistic model is accurate, then the SSM rates derived from the model should also be trustworthy. For this purpose, we compare the model-derived MSM rates against the GMM-Demux-classifier-observed MSM rates. If the numbers match, then we claim the GEM formation model must accurately characterizes the GEM formation process.

For comprehensiveness, we compare not only the overall MSM rates of a dataset, but also the MSM rates of individual sample combinations. For each sample combination, we compare the model-derived MSM UMI count against the MSM-classifier-observed UMI count. The comparison results are summarized into Venn diagrams, which illustrate the number of SSDs of each sample as well as the number of MSMs of each sample combination. We compare the model-derived Venn diagram against the MSM-classifier-observed Venn diagram. Figure 8 includes the Venn diagram comparisons of the PBMC-1 and the CD4^+^ Memory T datasets. Comparison of the PBMC-2 dataset is included in the Supplementary Materials (its per-sample-combination classification result cannot be visualized in a Venn diagram due to large number of sample combinations), in Table S8.

**Figure 8:**
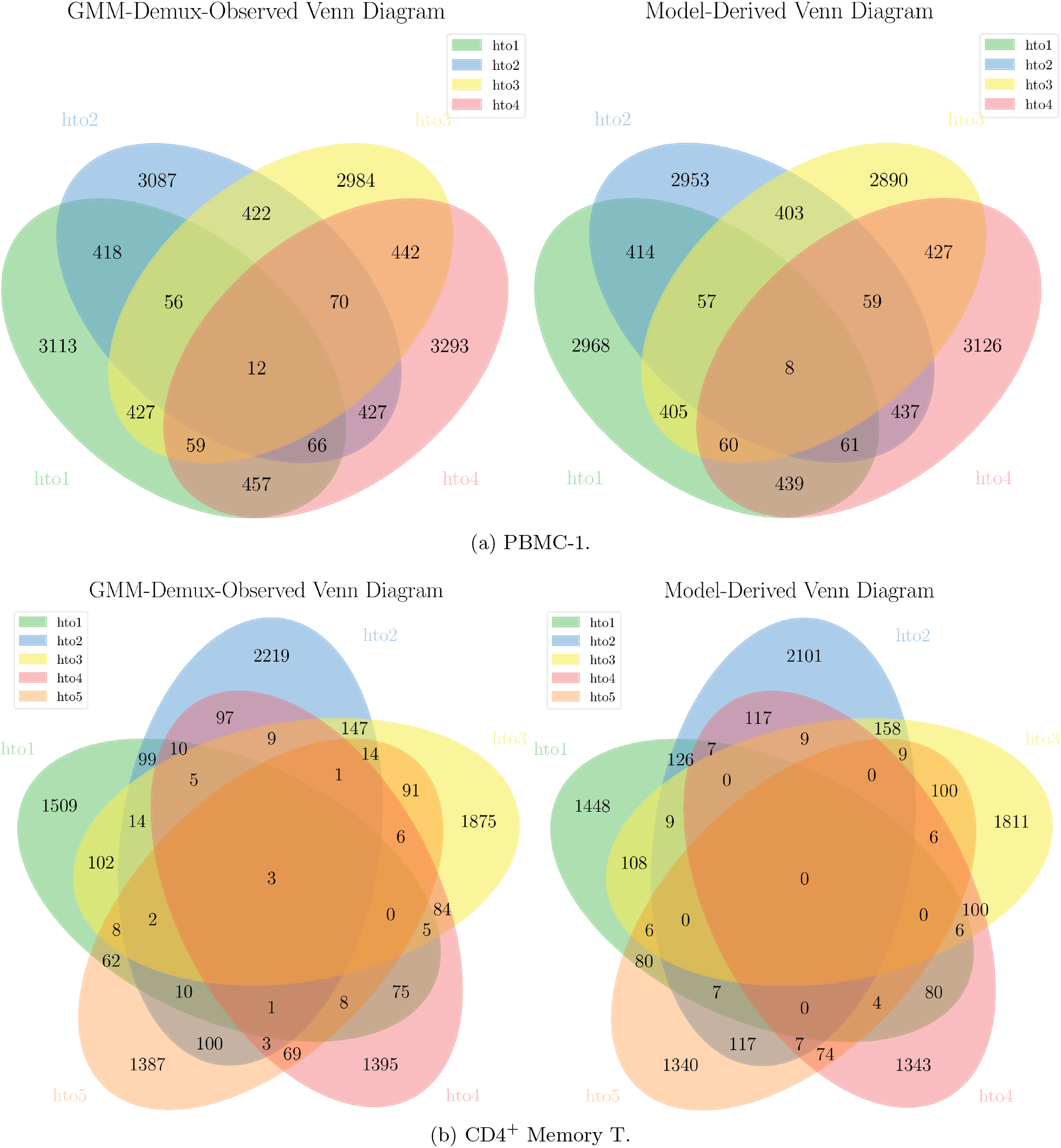
Comparison of model-derived Venn diagrams against GMM-Demux-observed Venn diagrams. Values in the model-derived Venn diagrams are consistent with values in the GMM-Demux-observed Venn diagrams, thus proving the correctness of the GEM-formation model.

From Figure 8 we observe that the model-derived MSM counts are consistent with the observed values from the MSM classifier. Therefore, we prove that the SSM rate estimator is accurate.

The estimated number of droplets (*X*); the model-estimated singlet, MSM (Est. MSM), SSM and relative SSM (RSSM) rates of each sample are summarized in Table 3. Also included in Table 3, are the GMM-Demux-classifier-observed MSM rates (Obs. MSM) and the proportions of unclear GEMs (GEMs with confidence scores below *c* = 0.8) and negative GEMs in each dataset^1^. Except the number of droplets (*X*), all rates are presented as percentiles (%). As shown in the table, the model-derived MSM rates are generally consistent with the classifier-observed MSM rates.

**Table 3:**
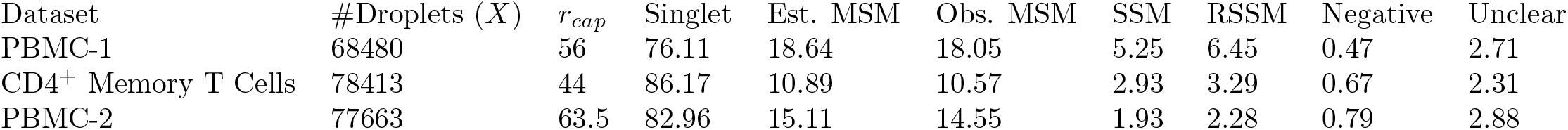
Summary of classification results across all datasets. All values except the number of droplets (*X*) are presented in percentages (%).

### 5.3 Online Experiment Planner Illustration

A screenshot of the online experiment planner is presented in Figure 9. To use the online experiment planner, the user provides the anticipated number of cells in the cell assay (*Y*), the number of HTO samples for sample barcoding (*M*), the estimated number of cell-assay droplets (*X*) and the estimated capture rate *r_cap_* of the single cell library prep equipment as inputs. The experiment planner outputs the estimated number of GEMs; the estimated singlet, MSM and SSM rates of GEM population; the estimated number of same-sample droplets (SSD) after removing MSMs using GMM-Demux; and the estimated RSSM rate of the remaining SSD population.

**Figure 9:**
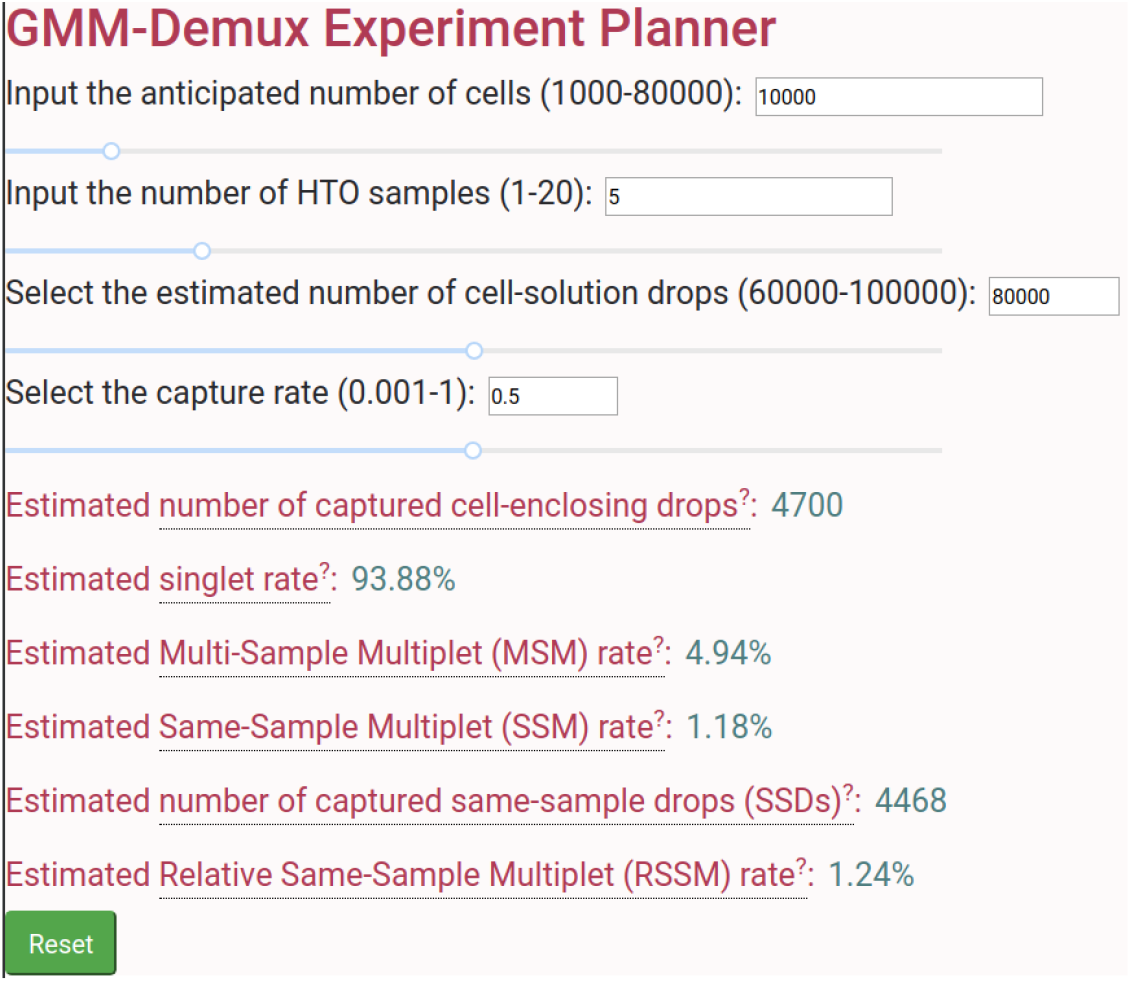
Screenshot of the online sample barcoding experiment planner. The experiment planner computes multiplet rates in real time as parameters are updated.

Both *X* and *r_cap_* can be obtained from the SSM rate estimator by running GMM-Demux over an existing sample barcoding scRNA-seq dataset. We observe that *X* typically ranges between 65K-80K and *r_cap_* typically varies between 0.4-0.6. In practice, we recommend profiling the library prep equipment by first conducting a pilot sample barcoding scRNA-seq experiment and

The online experiment planner dynamically interacts with the user. It dynamically updates the outputs upon changes on input values. Given a fixed number of cells as well as a fixed equipment profile (a fixed number of cell-assay droplets and a fixed capture rate), by dragging the slider handle of the HTO-sample-number input (*M*), the user can observe updates on multiplet rates in real time and find the minimum number of HTO samples required to meet a specific RSSM rate target.

### 5.4 Multiplet Rate Profiling Results

The profiles of the singlet, MSM, SSM rates, as well as the estimated number of harvested GEMs, under varying cell populations (*Y*) and sample counts (*M*) are shown in Figure 10. Figure 10 is generated under the setting of 80K cell-assay droplets (*X* = 80*K*) and 0.5 capture rate (*r_cap_* = 0.5). The above setting is based on the observation from Table 3.

**Figure 10:**
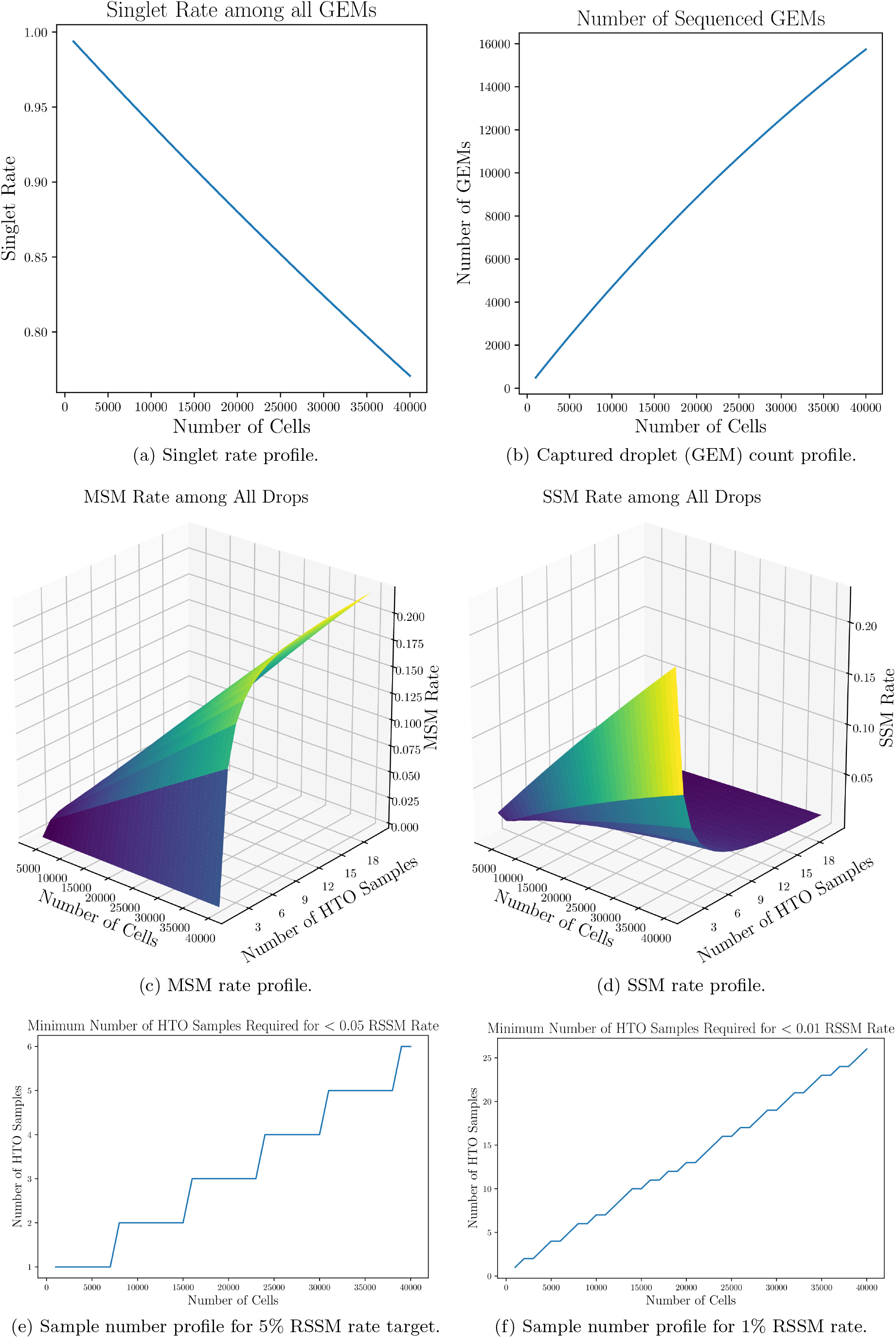
Sample barcoding experiment profiles under the assumption of 80K cell-assay droplets (*X* =80) and 0.5 capture rate (*r_cap_* = 50%). The singlet rate monotonically decreases as the cell population grows. Under a fixed cell population size, however, as the number of samples grows, more SSMs transit into MSMs.

As we have previously discussed, the singlet rate of an experiment, as well as the number of harvested gel-bead-enclosing GEMs only depend on the cell population size *Y* and are unaffected by the sample barcoding sample size *M*. Figure 10a and 10b displays the singlet rates and the estimated number of harvested GEMs under varying cell population sizes (*Y*). Between 1K to 40K, as more cells are loaded into the experiment, the singlet rate monotonically and super-linearly decreases, while the number of GEMs monotonically and sub-linearly increases.

The MSM and SSM rates, however, depend on both the cell population size *Y* and the sample number *M*. Under a fixed *Y*, a larger *M* transforms more SSMs into MSMs, although the margin quickly diminishes as *M* grows. On the other hand, under a fixed *M*, a larger *Y* leads to both greater MSM and greater SSM rates, in almost a linear fashion. The MSM and SSM rates under variate *Y*s and *M*s are shown in Figure 10c and 10d, respectively.

The singlet rates reported in Figure 10a are consistent with findings from previous works. As reported in Zheng *et al*. [45], the observed singlet rates are 98.4% for loading 2K cells and 96.9% for 6K cells, under the assumption of a 50% capture rate. In our work, according to the online experiment planner, under *X* = 80*K* and *r_cap_* = 0.5, the estimated singlet rate of *Y* = 2*K* is 98.75% while the singlet rate of *Y* = 6*K* is 96.3%.

Finally, we profile the minimum number of samples required to achieve certain RSSM rate targets under various cell population sizes (*Y*). Such profile helps in designing future sample barcoding experiments that minimize cost and complexity while meeting data quality targets. Figure 10f and 10e shows the profiling results of 1% and 5% RSSM rate targets, respectively, over various cell population sizes, again under the assumption of *X* = 80*K* and *r_cap_* = 0.5.

Singlet, MSM, SSM, RSSM rate profiles, as well as GEM count profiles under other settings ([*X, r_cap_*] other than [80*K*, 0.5]) are provided in the Supplementary Materials, from Figure S4 to S9. Together, they serve as guidance in designing future sample barcoding experiments.

### 5.5 Pure-Type GEM Verification Results

GEMs in the PBMC-1 dataset are manually assigned into 17 distinctive clusters following the gating strategy detailed in Maecker *et al*. [19], which is visualized in Figure 11. Among the 17 GEM clusters, 8 of them represent well-characterized cell types found in PBMCs (highlighted in green bounding boxes); 9 of them are rarely observed in PBMCs and are novel cell type candidates (highlighted in orange bounding boxes). All GEM clusters, annotated by their defining surface markers and their inferred cell types, if available, are summarized in Table 4.

**Figure 11:**
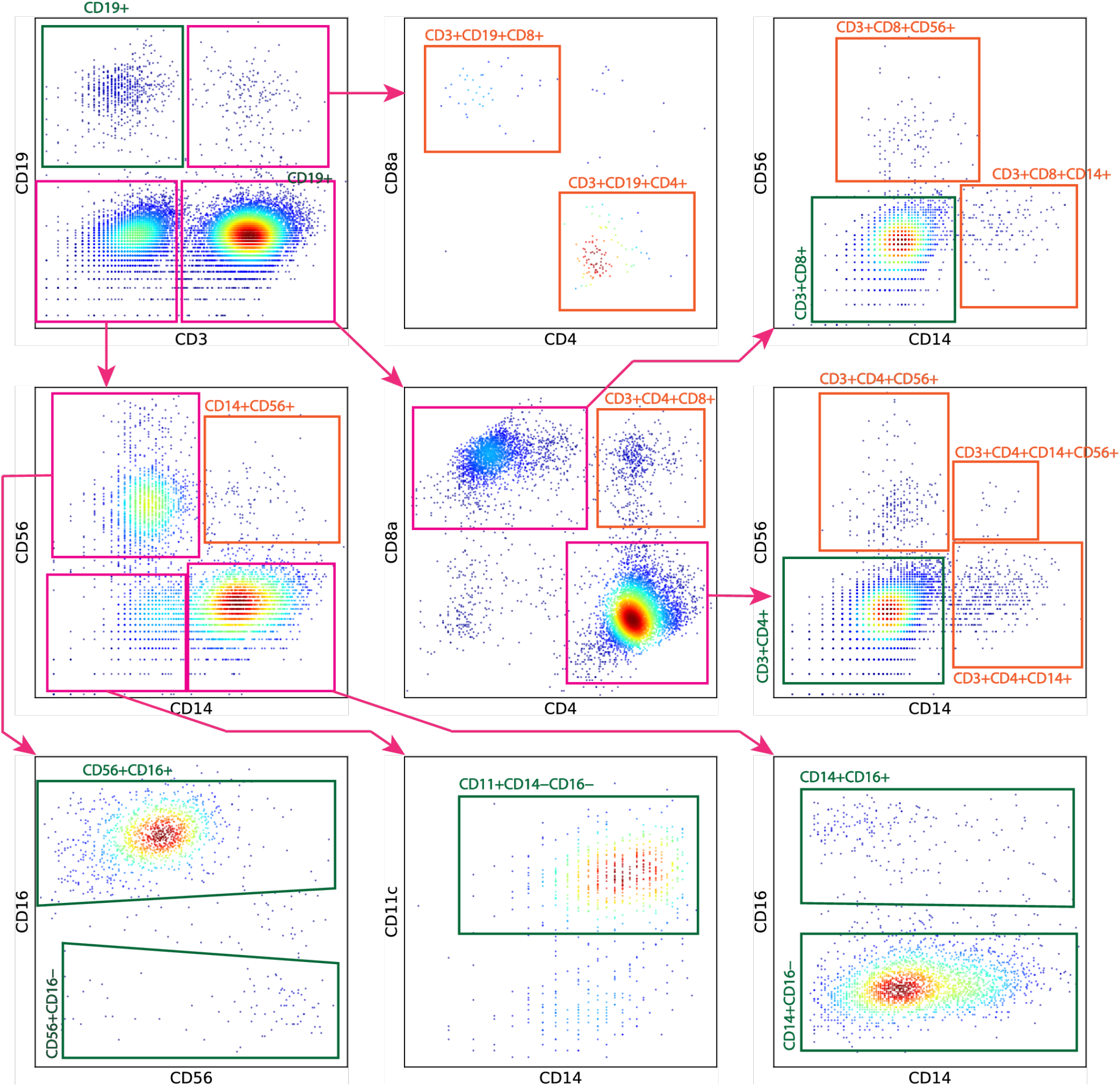
The manual gating strategy applied for cell type annotation in PBMC-1, using its surface marker expression data. In general, we follow the gating strategy outlined in Maecker *et al*. [19]. We first gate GEMs on CD3 and CD19. CD3^+^ GEMs are further gated over CD4 and CD8. CD3^−^CD19^−^ GEMs are further gated over CD14 and CD56. For GEM clusters gated from the CD14-CD56 penal, the CD14^+^ and CD56^+^ GEMs are further gated over CD16. The CD14^−^CD56^−^ GEMs are gated over CD11c to extract CD11c^+^ DC GEMs. The 8 cell type commonly observed in PBMCs are highlighted in green bounding boxes. Some GEM clusters, such as the CD3^+^CD4^+^ GEM cluster, go through additional gating, in order to reveal non-conventional sub-clusters in PBMCs (which are later classified as phony-type GEM clusters), such as the CD3^+^CD4^+^CD14^+^ GEM cluster or the CD3^+^CD4^+^CD56^+^ GEM cluster. Non-conventional GEM clusters are highlighted in orange bounding boxes.

**Table 4:** Summary of the 17 GEM clusters manually gated from PBMC-1. Given the cell hashing configuration, the minimum MSM percentage of a phony-type GEM cluster in PBMC-1 is 74.98%. Pure-type GEM clusters have variate MSM rates depending on their size. Among the 17 manually-gated GEM clusters, 9 have MSM percentages approaching and exceeding 74.98% and are classified as phony-type GEM clusters. 6 have MSM percentages of pure-type GEM clusters and are classified pure-type GEM clusters. 2 have MSM percentages of neither pure-type nor phony-type GEM clusters and are classified as mixture clusters.

| Cell Type | MSM %<br>(Observed) | MSM %<br>(Phony) | MSM %<br>(Pure) | p-value<br>(Phony) | p-value<br>(Pure) | Cluster<br>classification |
| --- | --- | --- | --- | --- | --- | --- |
| CD19<sup>+</sup> (B cells) | 6.47% | 74.98% | 5.93% | 0 | 0.29 | Pure |
| CD3<sup>+</sup>CD4<sup>+</sup> (Helper T cells) | 11.38% | 74.98% | 11.36% | 0 | 0.49 | Pure |
| CD3<sup>+</sup>CD8<sup>+</sup> (Cytotoxic T cells) | 7.33% | 74.98% | 7.52% | 0 | 0.68 | Pure |
| CD14<sup>+</sup>CD16<sup>-</sup> (Classical monocytes) | 7.00% | 74.98% | 7.49% | 0 | 0.88 | Pure |
| CD14<sup>+</sup>CD16<sup>+</sup> (Non-classical monocytes) | 14.91% | 74.98% | 5.82% | 1.75e-175 | 2.26e-13 | Mixture |
| CD56<sup>+</sup>CD16<sup>-</sup> (CD16<sup>-</sup> NK cells) | 6.20% | 74.98% | 6.41% | 8.70e-113 | 0.64 | Pure |
| CD56<sup>+</sup>CD16<sup>+</sup> (CD16<sup>+</sup> NK cells) | 9.62% | 74.98% | 6.72% | 0 | 1.00e-07 | Mixture |
| CD11<sup>+</sup>CD14<sup>-</sup>CD16<sup>-</sup> (DCs) | 7.30% | 74.98% | 5.86% | 1.01e-149 | 0.20 | Pure |
| CD14<sup>+</sup>CD56<sup>+</sup> | 76.56% | 74.98% | 6.90% | 0.62 | 5.24e-77 | Phony |
| CD3<sup>+</sup>CD4<sup>+</sup>CD14<sup>+</sup> | 74.67% | 74.98% | 6.22% | 0.42 | 0 | Phony |
| CD3<sup>+</sup>CD4<sup>+</sup>CD19<sup>+</sup> | 74.04% | 74.98% | 6.88% | 0.41 | 3.44e-119 | Phony |
| CD3<sup>+</sup>CD4<sup>+</sup>CD56<sup>+</sup> | 73.38% | 74.98% | 5.73% | 0.21 | 0 | Phony |
| CD3<sup>+</sup>CD4<sup>+</sup>CD8<sup>+</sup> | 75.55% | 74.98% | 6.02% | 0.66 | 0 | Phony |
| CD3<sup>+</sup>CD8<sup>+</sup>CD14<sup>+</sup> | 73.24% | 74.98% | 6.23% | 0.27 | 1.18e-165 | Phony |
| CD3<sup>+</sup>CD8<sup>+</sup>CD19<sup>+</sup> | 73.81% | 74.98% | 9.21% | 0.44 | 1.64e-41 | Phony |
| CD3<sup>+</sup>CD8<sup>+</sup>CD56<sup>+</sup> | 75.47% | 74.98% | 8.33% | 0.54 | 6.64e-57 | Phony |
| CD3<sup>+</sup>CD4<sup>+</sup>CD14<sup>+</sup>CD56<sup>+</sup> | 84.62% | 74.98% | 13.04% | 0.86 | 8.13e-14 | Phony |

For each GEM cluster, GMM-Demux computes the MSM percentage of the cluster and compares it against the anticipated pure-type MSM percentage as well as the anticipated phony-type MSM percentage of the cluster. The anticipated pure-type MSM percentage of the cluster is a hypothetical value derived from the GEM formation model by assuming that the GEM cluster is a pure-type GEM cluster. Similarly, the anticipated phony-type GEM percentage is computed by assuming the GEM cluster is a phony-type GEM cluster. Based on the observed and anticipated MSM percentages, GMM-Demux performs pure-type and phony-type hypothesis testings and classifies the GEM cluster according to the p-values of both tests. The classification results, as well as the intermediate results in classifying each GEM cluster, are also included in Table 4.

As summarized in Table 4, the PBMC-1 dataset contains 9 cell types rarely-observed in PBMCs. Named after their defining surface markers, these are: CD14^+^CD56^+^, CD3^+^CD4^+^CD14^+^, CD3^+^CD4^+^CD19^+^, CD3^+^CD4^+^CD56^+^, CD3^+^CD4^+^CD8^+^, CD3^+^CD8^+^CD14^+^, CD3^+^CD8^+^CD19^+^, CD3^+^CD8^+^CD56^+^ and CD3^+^CD4^+^CD14^+^CD56^+^. Upon further investigation, we observe that all 9 GEM clusters have very high MSM percentages, approaching and exceeding their anticipated phony-type MSM percentages. When tested with pure-type hypothesis, all 9 cluster have extremely small p-values; and large p-values from phony-type hypothesis tests. Consequently, GMM-Demux designates all 9 GEM clusters as phony-type clusters.

Such result suggests that all 9 GEM clusters contain multiplets of different cell types. For instance, the CD14^+^CD56^+^ GEM cluster contains multiplets that include both monocyte cells (CD14^+^) and NK cells (CD56^+^). Among the 9 phony-type GEM clusters, the CD3^+^CD4^+^CD14^+^CD56^+^ GEM cluster has the largest MSM percentage, significantly larger than the rest. With further examination of its defining surface markers, we conclude that it contains triple-type GEMs—GEMs that include CD3^+^CD4^+^ T cells, CD14^+^ monocytes and CD56^+^ NK cells. According to the GEM formation model for phony-type hypothesis testing, detailed in the Supplementary Materials section 3, triple-type phony GEM clusters have higher MSM percentages than double-type phony-type GEM clusters. This explains the larger MSM percentage of the CD3^+^CD4^+^CD14^+^CD56^+^ GEM cluster.

For the remaining 8 GEM clusters, which represent well-characterized cell types in PBMCs, 6 of them are classified as pure-type GEM clusters, with the exception of the CD14^+^CD16^+^ non-classical monocyte GEM cluster and the CD56^+^CD16^+^NK GEM cluster. Both clusters are classified as mixture GEM clusters, suggesting that they contain both pure-type and phony-type GEMs. This classification result is reasonable, as both GEM clusters contain fractions of indistinguishable multiplets. For instance, inside the CD14^+^CD16^+^ GEM cluster, there could be a small fraction of CD14^+^CD16^+^-and-CD14^+^CD16^−^ phony-type GEMs. These phony-type GEMs are indistinguishable from the CD14^+^CD16^+^ pure-type GEMs in gating. In gating, boundaries between cell types are drawn in a log-transformed surface marker space. After log transformation, the surface marker expression profile of a CD14^+^CD16^+^-and-CD14^+^CD16^−^ phony-type GEM is almost identical to a CD14^+^CD16^+^ pure-type GEM, even if they contain the same CD14^+^CD16^+^ non-classical monocyte cell. The only difference: the CD14^+^CD16^+^-and-CD14^+^CD16^−^ phony-type GEM is likely to have a slightly larger log-transformed CD14 expression value. Such subtle differences do not warrant the separation of CD14^+^CD16^+^-and-CD14^+^CD16^−^ phony-type GEMs from CD14^+^CD16^+^ pure-type GEMs. Due to intrinsic variations in surface marker expression levels, the two types of GEMs intermix with each other into a single, indivisible GEM cluster. Similarly, CD56^+^CD16^+^-and-CD56^+^CD16^−^ phonytype GEMs are also indistinguishable from CD56^+^CD16^+^ pure-type GEMs. This explains the slightly-higher-than-expected MSM percentages in the CD14^+^CD16^+^ monocyte GEM cluster and the CD56^+^CD16^+^ NK GEM cluster, which resulted in designating them as mixture GEM clusters. Nonetheless, these should be the only phony-type GEMs they contain. Therefore, the MSM percentages of both clusters are only moderately above their corresponding pure-type MSM percentages, remaining significantly smaller than their corresponding phony-type MSM percentages, reflecting that both clusters still have a pure-type GEM majority. Overall, we conclude that these 8 GEM clusters represent real cell types in PBMC, in concordance with previous knowledge on PBMCs[19].

To validate the classification results of GMM-Demux, we conducted an additional CITE-seq sequencing experiment over a PBMC sample from the same donor of PBMC-1. The additional CITE-seq experiment measures the same set of surface markers as in PBMC-1. To control the percentage of multiplets, we loaded only 3.2K cells while harvesting ~1.6K GEMs. The estimated percentage of multiplets of this dataset is 1.9%, compared to 23.9% in PBMC-1. We sorted GEMs following the same gating strategy illustrated in Figure 11. Table 5 records the percentages of the 17 manually-gated cell types in both PBMC-1 and the validation dataset. We observe that all 9 phony-type GEM clusters identified in PBMC-1 have much-reduced, close-to-zero presences in the validation dataset, while the 8 cell-type-defining GEM clusters have similar footprints. This confirms the classification results of GMM-Demux.

**Table 5:**
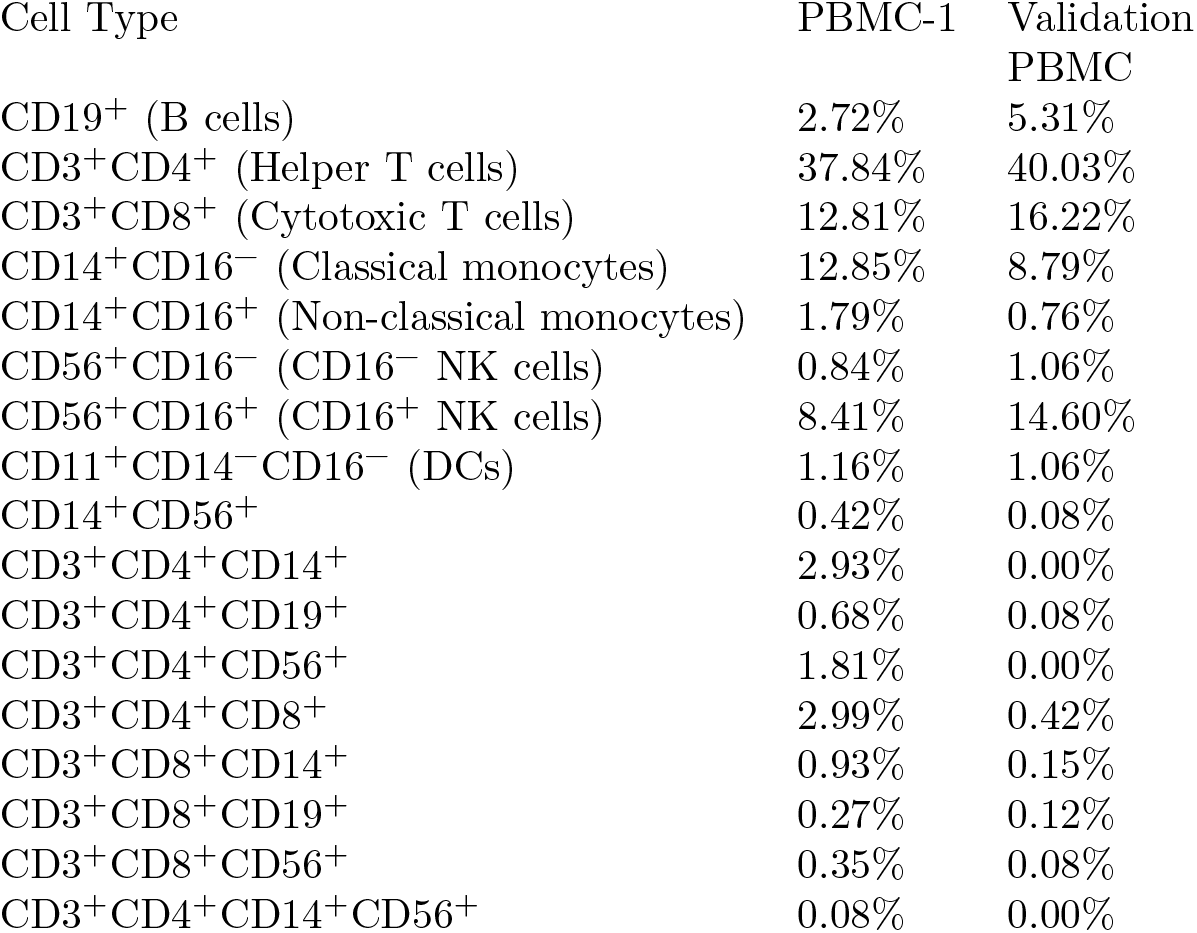
Percentages of the 17 cell type in both PBMC-1 and the validation dataset. All phony-type GEM clusters identified in PBMC-1 have close-to-zero presences in the validation dataset, suggesting that these cell types do not really exist and are artifacts of multiplets.

| Cell Type | PBMC-1 | Validation<br>PBMC |
| --- | --- | --- |
| $CD19^+$ (B cells) | 2.72% | 5.31% |
| $CD3^+CD4^+$ (Helper T cells) | 37.84% | 40.03% |
| $CD3^+CD8^+$ (Cytotoxic T cells) | 12.81% | 16.22% |
| $CD14^+CD16^-$ (Classical monocytes) | 12.85% | 8.79% |
| $CD14^+CD16^+$ (Non-classical monocytes) | 1.79% | 0.76% |
| $CD56^+CD16^-$ ($CD16^-$ NK cells) | 0.84% | 1.06% |
| $CD56^+CD16^+$ ($CD16^+$ NK cells) | 8.41% | 14.60% |
| $CD11^+CD14^-CD16^-$ (DCs) | 1.16% | 1.06% |
| $CD14^+CD56^+$ | 0.42% | 0.08% |
| $CD3^+CD4^+CD14^+$ | 2.93% | 0.00% |
| $CD3^+CD4^+CD19^+$ | 0.68% | 0.08% |
| $CD3^+CD4^+CD56^+$ | 1.81% | 0.00% |
| $CD3^+CD4^+CD8^+$ | 2.99% | 0.42% |
| $CD3^+CD8^+CD14^+$ | 0.93% | 0.15% |
| $CD3^+CD8^+CD19^+$ | 0.27% | 0.12% |
| $CD3^+CD8^+CD56^+$ | 0.35% | 0.08% |
| $CD3^+CD4^+CD14^+CD56^+$ | 0.08% | 0.00% |

Table 5 proves that removing MSMs alone does not eliminate all multiplets. None of the phony GEM clusters has a MSM percentage of 100%. All phony GEM clusters have non-negligible fractions of SSMs, which cannot be revealed or removed through sample barcoding alone. After removing all phony-type GEM clusters, we estimate the RSSM of PBMC-1 is further reduced to 3.29%, from 6.45%.

## 6 Conclusion and Discussion

In this paper, we proposed a model-based Bayesian framework, GMM-Demux, for detecting sample-barcoding-detectable multiplets in a sample barcoding dataset; estimating the percentage of sample-barcoding-undetectable multiplets in the remaining dataset; estimating the multiplet rates of a planned future sample barcoding experiment; and validating the existence of specific cell types. At its core, GMM-Demux uses Gaussian-mixture models to identify GEMs that contain sample-specific cells and then uncovers MSMs by selecting GEMs that contain cells from multiple samples. We showed that GMM-Demux accurately and consistently classifies GEMs into SSDs and MSMs and generates more accurate and more consistent results when compared against existing methods. We further proposed a GEM formation model to estimate the SSM rate in a sample barcoding dataset. The GEM formation model describes the GEM formation process as an augmented binomial process. We showed that the GEM formation model accurately characterizes the GEM formation process. We built an online experiment planner that estimates the multiplet rate of a planned sample barcoding (or an ordinary single cell) experiment. Then we used the online experiment planner to generate a series of multiplet profiles under various experimental setups. Finally, we proposed a GEM cluster classifier that checks the existence of cell-type defining GEM clusters, and showed that GMM-Demux correctly identifies phony-type GEM clusters in single cell datasets.

GMM-Demux is the first work that is able to not only accurately and consistently classify sample-barcoding-detectable MSMs in a sample barcoding dataset, but also estimate the undetectable SSM rates among the remaining SSDs. Furthermore, GMM-Demux is the first work attempting to model the GEM formation process using a generative model. GMM-Demux incorporates its GEM formation model into an online experiment planner that is capable of anticipating experimental outcomes of planned sample barcoding experiments; and it is a first in systematically verifying the legitimacy of putative cell types in sample barcoding.

Despite out-performing existing methods, the underlying assumptions of GMM-Demux impose a number of limitations. First, GMM-Demux assumes a wide gap exists in the HTO concentrations before and after pooling samples. Such differences are key to defining the two peaks in the bimodal distribution of HTO UMI counts. Although from our observation, the two peaks are always well defined and are always far apart from each other in the HTO UMI count distributions, this is not 100% guaranteed, especially when the sample number is low (e.g., *M* = 2). When pooling fewer samples together, the HTO concentration reduction by pooling could diminish. However, this is more of a limitation of the sample barcoding technology, rather than a limitation of GMM-Demux. Based on the premise of the sample barcoding technology, which strives to tag only sample-specific cells with HTOs, we believe that the bimodal distribution assumption should always hold. Second, the online experiment planner requires prior knowledge of the number of cell-assay droplets generated by the library prep equipment. We suggest that users should profile their library prep equipment once with GMM-Demux for the cell-assay droplet count and use the profiled number in future experiment planning. While it is logical to assume that the same library prep equipment generates the same number of cell-assay droplets over repetitive runs, this is yet to be confirmed. In reality, based on the total volume of the loaded cell assay, the total count of cell-assay droplets could vary, even if the cell assay pump operates at a constant frequency. Such variation however, does not affect the MSM classifier or the SSM rate estimator and it can be potentially alleviated by maintaining a series of profiles for variate cell assay volumes. Finally, GMM-Demux assumes cells are partitioned into droplets independently. This model does not consider the volume taken up by each cell. A more realistic model would assign diminishing likelihoods to having additional cells partitioned into a droplet as more cells accumulate in the droplet. To that end, GMM-Demux does not take cell-size differences into consideration either. As cells differ in sizes, a more accurate model would assign a smaller likelihood to having two large cells partitioned into the same droplet than that of two small cells. Unfortunately, the cell size and droplet size information is not readily available in sample barcoding data, which limits us from studying the effect of cell size on multiplet rates. Nevertheless, given that the probability of a droplet containing more than three cells is already close to zero according to our current droplet formation model, and the fact that the cell-assay droplet size has to be large enough to accommodate the largest possible cell in a tissue, we believe it is unnecessary to further complicate the GEM formation model to include the cell size information.

In our future work, we intend to perform more sample barcoding experiments with different tissues and investigate the underlying mechanisms that govern the number of cell-assay droplets and the capture rate in a sample barcoding experiment. We seek to expand the GEM formation model and use it to detect false lineage discoveries and false cell type discoveries in single cell data analysis. We also plan to investigate how to identify SSMs within SSDs.

## Supporting information

Supplementary sections

## 7 Acknowledgements

This work is supported by National Institute of Health grants R01HL137709 (W.C. and K.C.), U01DK062420 (W.C. and R.D.), and Childrens Hospital of Pittsburgh (W.C., R.D., Y.J., Q.L, Q.Y. and H.X.).

## 8 Code Availability

GMM-Demux is implemented in Python using the SciPy library[11]. The online sample barcoding experiment planner is built using the Dash framework[30].

The source code of GMM-Demux is freely available as at https://github.com/CHPGenetics/GMM-demux. The online experiment planner is publicly accessible at http://www.pitt.edu/~wec47/gmmdemux.html A detailed tutorial is also included in the above repository.

## 9 Data Availability

Data will be made publicly available upon acceptance of the paper.

## Footnotes

1 Singlet, MSM, SSM and RSSM rates, as well as the droplet number *X* are computed after the removal of negative and unclear GEMs. Negative and unclear rates, however, are computed with regard to the entire GEM population

