## Supplementary sections for "Sample demultiplexing, multiplet detection, experiment planning and novel cell type verification in single cell sequencing"

### Supplementary Materials

#### 1 Modeling the General Droplet Formation Process

Outlined in the original droplet-seq paper[18], in droplet-based single cell library prep, GEMs are formed by "co-flowing" two aqueous solutions across an oil channel to form nanoliter-sized droplets. One flow contains gel beads, microcapsules that contain barcoded microparticles suspended in a lysis buffer. The other flow contains the cell assay, which is a HTO-tagged cell suspension. Figure 1 illustrates the co-flow process. In our model, the two flows are driven by two separate pumps that operate in discrete pulses. Each pulse pumps unit volume solution into oil, forming an emulsion droplet. In our model, the two pumps does not operate in sync and their frequencies differ. Assuming the unit volume of each pulse from the cell assay pump is $\mu$, and the total volume of the cell assay solution is $V$, then the total number of pulses from the cell-assay pump throughout a sample barcoding library prep can be calculated as $X = \frac{V}{\mu}$. Therefore, in the final solution, there are a total of $X = \frac{V}{\mu}$ droplets that contain cell-assay solution.

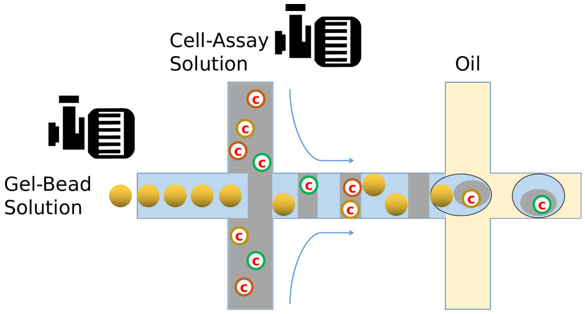


Figure 1: Illustration of the co-flow process.

Not all cell-assay droplets can be recovered through sequencing. A droplet is sequenced only if it also contains a gel bead, which is not always guaranteed. In our model, we assume $r_{cap}$ as a constant probability of a droplet getting a gel bead. Cell-assay droplets without gel beads are not harvested through sequencing.

#### 2 Validating Multiplet Rate Equations

We validate the multiplet rate equations by comparing the equation-derived multiplet rates against the multiplet rates obtained through simulation. Specifically, given $[Y, M, X]$ ($Y$ is the total number of cells, $M$ is the number of samples and $X$ is the total number of cell-assay droplets), we divide $Y$ cells evenly into $M$ samples and randomly partition each cell into a droplet. We do not include $r_{cap}$ in our simulation as $r_{cap}$ does not affect the multiplet rates. We collect all droplets from the simulation; count the numbers of singlets, SSMs and MSMs and computes the singlet, SSM and MSM rates. For each $[Y, M, X]$ parameter set, we repeat the simulation 500, 1000 and 1500 times and average the singlet, SSM and MSM rates over all iterations. The comparison results are shown in Table 1, 2 and 3.

As shown in the tables, as the number of iterations increases, the multiplet rates obtained through simulation asymptotically approaches the model-derived multiplet rates. This suggests that the equations for computing multiplet rates are correct.

| #Cells ($Y$) | #Droplets ($X$) | #Samples ($M$) | 500 iterations | 1000 iterations | 1500 iterations | model-derived |
|---|---|---|---|---|---|---|
| 2000 | 8000 | 2 | 0.88038 | 0.88029 | 0.88040 | 0.88025 |
| 2000 | 8000 | 5 | 0.88001 | 0.88001 | 0.88030 | 0.88025 |
| 2000 | 8000 | 10 | 0.88042 | 0.88003 | 0.88028 | 0.88025 |
| 2000 | 10000 | 2 | 0.90313 | 0.90296 | 0.90349 | 0.90337 |
| 2000 | 10000 | 5 | 0.90278 | 0.90301 | 0.90344 | 0.90337 |
| 2000 | 10000 | 10 | 0.90332 | 0.90364 | 0.90337 | 0.90337 |
| 4000 | 8000 | 2 | 0.77068 | 0.77084 | 0.77069 | 0.77078 |
| 4000 | 8000 | 5 | 0.77029 | 0.77085 | 0.77129 | 0.77078 |
| 4000 | 8000 | 10 | 0.77041 | 0.77062 | 0.77078 | 0.77078 |
| 4000 | 10000 | 2 | 0.81304 | 0.81327 | 0.81298 | 0.81332 |
| 4000 | 10000 | 5 | 0.81287 | 0.81330 | 0.81353 | 0.81332 |
| 4000 | 10000 | 10 | 0.81403 | 0.81362 | 0.81349 | 0.81332 |

Table 1: Comparison of simulated and model-derived singlet rates.

| #Cells ($Y$) | #Droplets ($X$) | #Samples ($M$) | 500 iterations | 1000 iterations | 1500 iterations | model-derived |
|---|---|---|---|---|---|---|
| 2000 | 8000 | 2 | 0.06238 | 0.06258 | 0.06225 | 0.06242 |
| 2000 | 8000 | 5 | 0.09773 | 0.09758 | 0.09736 | 0.09742 |
| 2000 | 8000 | 10 | 0.10842 | 0.10890 | 0.10860 | 0.10870 |
| 2000 | 10000 | 2 | 0.05042 | 0.05014 | 0.04988 | 0.04996 |
| 2000 | 10000 | 5 | 0.07895 | 0.07869 | 0.07833 | 0.07836 |
| 2000 | 10000 | 10 | 0.08761 | 0.08728 | 0.08751 | 0.08757 |
| 4000 | 8000 | 2 | 0.12455 | 0.12417 | 0.12434 | 0.12436 |
| 4000 | 8000 | 5 | 0.18978 | 0.18924 | 0.18887 | 0.18940 |
| 4000 | 8000 | 10 | 0.21019 | 0.20976 | 0.20957 | 0.20967 |
| 4000 | 10000 | 2 | 0.10001 | 0.09959 | 0.09996 | 0.09967 |
| 4000 | 10000 | 5 | 0.15366 | 0.15332 | 0.15321 | 0.15329 |
| 4000 | 10000 | 10 | 0.16937 | 0.16994 | 0.17004 | 0.17022 |

Table 2: Comparison of simulated and model-derived MSM rates.

| #Cells ($Y$) | #Droplets ($X$) | #Samples ($M$) | 500 iterations | 1000 iterations | 1500 iterations | model-derived |
|---|---|---|---|---|---|---|
| 2000 | 8000 | 2 | 0.05723 | 0.05712 | 0.05733 | 0.05732 |
| 2000 | 8000 | 5 | 0.02225 | 0.02239 | 0.02233 | 0.02232 |
| 2000 | 8000 | 10 | 0.01114 | 0.01105 | 0.01111 | 0.01104 |
| 2000 | 10000 | 2 | 0.04644 | 0.04688 | 0.04662 | 0.04666 |
| 2000 | 10000 | 5 | 0.01825 | 0.01828 | 0.01822 | 0.01826 |
| 2000 | 10000 | 10 | 0.00906 | 0.00907 | 0.00910 | 0.00904 |
| 4000 | 8000 | 2 | 0.10475 | 0.10497 | 0.10496 | 0.10485 |
| 4000 | 8000 | 5 | 0.03992 | 0.03990 | 0.03982 | 0.03980 |
| 4000 | 8000 | 10 | 0.01938 | 0.01961 | 0.01964 | 0.01954 |
| 4000 | 10000 | 2 | 0.08693 | 0.08712 | 0.08705 | 0.08699 |
| 4000 | 10000 | 5 | 0.03345 | 0.03336 | 0.03325 | 0.03337 |
| 4000 | 10000 | 10 | 0.01658 | 0.01643 | 0.01645 | 0.01644 |

Table 3: Comparison of simulated and model-derived SSM rates.

### 3 Hypothesis Testings in Pure-Type GEM Verification

To classify a GEM cluster $G$, GMM-Demux tests two hypotheses: $G$ is a pure-type GEM cluster; and $G$ is a phony-type GEM cluster. In both hypothesis tests, GMM-Demux first computes the expected value of $P(i \in \texttt{MSM}_\texttt{G}|i \in G)$—the probability of a GEM in $G$ being a MSM, under the respective hypotheses. Then GMM-Demux computes the statistical significance of observing the true MSM population of $G$ under the expected $P(i \in \texttt{MSM}_\texttt{G}|i \in G)$ derived from the previous step. GMM-Demux tests both hypotheses with the binomial test: let $P_0(i \in \texttt{MSM}_\texttt{G}|i \in G)$ denote the expected value of $P(i \in \texttt{MSM}_\texttt{G}|i \in G)$; GMM-Demux assumes that each GEM $i$ in $G$ has the probability of $P_0(i \in \texttt{MSM}_\texttt{G}|i \in G)$ to become a MSM, and computes the statistical significance (the p-value) of observing the true number of MSMs in $G$. The null hypotheses of the binomial test is rejected if the p-value of the test falls below a user defined alpha level (0.05 by default). According to the binomial test results of both hypotheses, $G$ is classified as a pure-type GEM cluster, a phony-type GEM cluster or a mixture cluster accordingly.

GMM-Demux derives $P_0(i \in \texttt{MSM}_\texttt{G}|i \in G)$ from the GEM formation model. For simplicity, we assume cells are uniformly randomly distributed into $M$ sample barcoding samples. Formally, the probability of a GEM $i$ in $G$ is a MSM can be formulated as the following:

$$P_0(i \in \texttt{MSM}_\texttt{G}|i \in G) = \sum_j P(i \in \texttt{MSM}_\texttt{G}|c_i = j) \cdot P(c_i = j) \tag{20}$$

where $c_i$ denotes the number of cells in GEM $i$ and $c_i = j$ states that there are $j$ cells in $i$. If $G$ is a phony-type cluster, since all GEMs are multiplets, $P(c_i = 1) = 0$. For $j > 1$, $P(i \in \texttt{MSM}_\texttt{G}|c_i = j) = 1 - P(i \in \texttt{SSM}_\texttt{G}|c_i = j) = 1 - \binom{1}{M}(\frac{1}{M})^j = 1 - (\frac{1}{M})^{j-1}$. Let $k$ denote the number of cell types in a GEM. If $G$ is a $k$-cell-type phony-type cluster, where each GEM in $G$ contains (the same) $k$ ($k > 1$) cell types, then $P_0(i \in \texttt{MSM}_\texttt{G}|i \in G) > 1 - (\frac{1}{M})^{k-1}$, as $P(c_i = j) = 0$ for all $j < k$. In general, let $y_G$ represent the total cell population size of the $k$ cell types of $G$ in the cell assay. In a $k$-cell-type phony-type cluster, $P(c_i = j) = 0$ for all $j < k$ and $P(c_i = j) = \binom{y_G - k}{j - k}(\frac{1}{X})^{j-k}(1 - \frac{1}{X})^{k-j} \cdot P(c_i = k)$ for all $j \geq k$. When $X > y_G >> 1$ and $k > 1$, we have $P_0(i \in \texttt{MSM}_\texttt{G}|i \in G) \gtrapprox 1 - (\frac{1}{M})^{k-1}$.

The formula for $P_0(i \in \texttt{MSM}_\texttt{G}|c_i = j)$ is slightly different in more complex scenarios where the cell assay is not evenly distributed into the $M$ sample barcoding samples. Assume cells are not uniformly randomly distributed into sample barcoding samples and let $p_x$ denote the probability of a cell being distributed to a sample $x$. Then $P(i \in \texttt{SSM}_\texttt{G}|c_i = j) = \sum_{x=1}^{M} P(i \in \texttt{SSM}_\texttt{G}{}^x|c_i = j)$, where $P(i \in \texttt{SSM}_\texttt{G}{}^x|c_i = j)$ denotes the probability of a droplet $i$ being a SSM of sample $x$, given that $i$ already contains $j$ cells. In other words, $P(i \in \texttt{SSM}_\texttt{G}{}^x|c_i = j)$ is the probability of all $j$ cells in $i$ come from sample $x$, which means $P(i \in \texttt{SSM}_\texttt{G}{}^x|c_i = j) = (p_x)^j$. Therefore, we have $P(i \in \texttt{SSM}_\texttt{G}|c_i = j) = \sum_{x=1}^{M}(p_x)^j$. Accordingly, $P_0(i \in \texttt{MSM}_\texttt{G}|i \in G)$ is updated as $P_0(i \in \texttt{MSM}_\texttt{G}|i \in G) \gtrapprox 1 - \sum_{x=1}^{M}(p_x)^k$ under the condition of $X > y_G >> 1$ and $k > 1$.

With $P_0(i \in \texttt{MSM}_\texttt{G}|c_i = j)$ computed for $k > 1$, the null and alternative hypotheses of the phony-type hypothesis test are formulated as follows: the null hypothesis states that $H_0^{phony} : P(i \in \texttt{MSM}_\texttt{G}|i \in G) \geq P_0(i \in \texttt{MSM}_\texttt{G}|i \in G)$, while the alternative hypothesis states that $H_a^{phony} : P(i \in \texttt{MSM}_\texttt{G}|i \in G) < P_0(i \in \texttt{MSM}_\texttt{G}|i \in G)$. The null hypothesis is tested with the binomial test and is rejected if the p-value of the test is smaller than the alpha level. Rejecting the null hypothesis suggests that $G$ is not a phony-type GEM cluster.

The phony-type hypothesis test can be further extended to infer the number of cell types ($k$) in $G$. For simplicity, again cells are assumed to be uniformly distributed into $M$ sample barcoding samples, without loss of generality. For $k = 2$, $P_0(i \in \texttt{MSM}_\texttt{G}|i \in G) \gtrapprox 1 - \frac{1}{M}$. For $k = 3$, $P_0(i \in \texttt{MSM}_\texttt{G}|i \in G) \gtrapprox 1 - (\frac{1}{M})^2$. For $k = n$, $P_0(i \in \texttt{MSM}_\texttt{G}|i \in G) \gtrapprox 1 - (\frac{1}{M})^{n-1}$. For each $P_0$, GMM-Demux performs the binomial test and rejects the null hypothesis when the p-value drops below the alpha value. Since $P_0(i \in \texttt{MSM}_\texttt{G}|i \in G)$ increases as $k$ increases, rejecting the null hypothesis suggests that $G$ is not a phony-type GEM cluster of no-less-than $k$ cell types (but could still be a phony-type GEM cluster with fewer cell types).

Rejecting $H_0^{phony}$ does not make $G$ a pure-type GEM cluster. $G$ could also be a mixture cluster, which contains both pure-type GEMs and phony-type GEMs. To check if $G$ is a pure-type GEM only cluster, GMM-Demux performs the *pure-type hypothesis test* $H^{pure}$, which assumes that $G$ includes only $k = 1$ cell type.

When $k = 1$, $P_0(i \in \texttt{MSM}_\texttt{G}|i \in G)$ is much smaller than $1 - \frac{1}{M}$. Let $\tau$ denote the cell type of $G$ and assume that $G$ includes all pure-type GEMs of type $\tau$. Let $y_\tau$ denote the number of $\tau$ cells in the cell assay and $z_G$ denote the number of GEMs in $G$. According to the GEM formation model, the probability of a droplet $i$ being a pure-type GEM of type $\tau$ can be computed as $P(i \in SSD_\tau) = (1 - (1 - \frac{1}{X})^{y_\tau})(1 - \frac{1}{X})^{Y - y_\tau}$. The probability of observing $z_G$ pure-type GEMs of type $\tau$ equals to $P(\#_{\texttt{GEM}_\tau} = z_G) = \binom{z_G/r_{cap}}{X} P(i \in SSD_\tau)^{z_G/r_{cap}}(1 - P(i \in Pure_\tau))^{X - (z_G/r_{cap})}$. Consequently, GMM-Demux generates an estimation of $y_\tau$ by maximizing the likelihood of $P(\#_{\texttt{GEM}_\tau} = z_G)$.

With $y_\tau$ computed, the probability of a droplet $i$ being a pure-type MSM of type $\tau$, denoted as $P(i \in \texttt{MSM}_\tau)$, can be computed in a manner that is similar to Equation 10. Subsequently, we have $P_0(i \in \texttt{MSM}_\texttt{G}|i \in G) = \frac{P(i \in \texttt{MSM}_\tau) \cdot X}{z_G}$. The null and alternative hypotheses of the pure-type hypothesis test are formulated as $H_0^{pure} : P_{i \in MSM_\tau} \leq P(i \in \texttt{MSM}_\tau)_0$ and $H_a^{pure} : P(i \in \texttt{MSM}_\tau) > P(i \in \texttt{MSM}_\tau)_0$, respectively. GMM-Demux tests the null hypothesis with the binomial test; computes the p-value; compares the p-value against the alpha level and rejects $H_0^{pure}$ if the p-value is smaller than the alpha level. Rejecting $H_0^{pure}$ suggests that $G$ is not a pure-type GEM cluster.

Based on the hypothesis testing results of both the pure-type and phony-type hypothesis tests, $G$ is classified as a pure-type GEM cluster, a phony-type GEM cluster or a mixture cluster. Specifically, $G$ is classified as a pure-type GEM cluster if

$H_0^{pure}$ is not rejected; $G$ is classified as a phony-type GEM cluster if $H_0^{phony}$ is not rejected; and $G$ is classified as a mixture cluster if both $H_0^{pure}$ and $H_0^{phony}$ are rejected.

$G$ has the potential to reveal a new cell type only if $G$ is classified as a pure-type cluster. When classified as a phony-type cluster, all GEMs in $G$ are recommended to be removed from the dataset, as all SSDs in $G$ has a high probability to be SSMs. Finally, being classified as a mixture cluster suggests that $G$ is not homogeneous and the clustering quality is low. Further refinement over $G$ is recommended.

### 4 Example GMM-Demux Output

GMM-Demux generates three output files. It generates a classification result file, a classification header file, and a summary file. An example result and header file set is presented in Table 4 and 5. In the classification result file, each sample and each multi-sample combination is given a unique classification id. The classification id is matched with a description text in the header file. The summary file contains two tables, illustrated in Table 6 and 7. The first table presents a summary of the computed and inferred parameters of the dataset, including the number of cell-assay droplets ($X$), the capture rate ($r_{cap}$), the singlet, MSM and SSM rates of the entire dataset. The second table contains a per-sample summary, which stores the estimated SSM rate of each sample.

| GEM Barcodes | Classification id | Confidence Score |
|---|---|---|
| AAACCTGAGACAAAGG-1 | 2 | 0.9999996 |
| AAACCTGAGACAAGCC-1 | 3 | 0.9999854 |
| AAACCTGAGAGGTTGC-1 | 3 | 0.9999995 |
| AAACCTGAGAGTGAGA-1 | 5 | 0.9952781 |
| AAACCTGAGCCCAACC-1 | 3 | 0.9999971 |
| AAACCTGAGCTTATCG-1 | 7 | 0.9999966 |
| AAACCTGAGGACAGAA-1 | 1 | 0.9999986 |
| AAACCTGAGGGATCTG-1 | 3 | 0.9903349 |
| AAACCTGAGGGATGGG-1 | 1 | 0.9995598 |
| AAACCTGAGGGCACTA-1 | 3 | 0.5391441 |
| AAACCTGAGGTTACCT-1 | 2 | 0.9999990 |
| AAACCTGAGTACGATA-1 | 2 | 0.9991886 |
| AAACCTGCAACACCCG-1 | 7 | 0.9998400 |
| AAACCTGCAAGCTGTT-1 | 3 | 0.9999598 |
| AAACCTGCAATCGAAA-1 | 0 | 0.9784425 |
| AAACCTGCACCCTATC-1 | 4 | 0.9995443 |

Table 4: An example MSM classification result file.

| Classification id | Classification Description |
|---|---|
| 0 | negative |
| 1 | hto1 |
| 2 | hto2 |
| 3 | hto3 |
| 4 | hto1-hto2 |
| 5 | hto1-hto3 |
| 6 | hto2-hto3 |
| 7 | hto1-hto2-hto3 |

Table 5: The header file of Table 4.

| #Droplets ($X$) | $r_{cap}$ | Est. Singlet | Obs. MSM | Est. SSM | Est. RSSM | Negative | Unclear |
|---|---|---|---|---|---|---|---|
| 68480 | 56 | 76.11 | 18.05 | 5.25 | 6.45 | 0.47 | 2.71 |

Table 6: An example summary file for the entire dataset. Except the number of droplets (X), all other values are in %.

| | hto1 | hto2 | hto3 | hto4 |
|---|---|---|---|---|
| # Cells | 8982 | 8937 | 8746 | 9462 |
| RSSM rate | 6.4141 | 6.3828 | 6.2494 | 6.7489 |

Table 7: The per-sample summary of the example sample barcoding experiment. All RSSM values are in %.

### 5 Distinct MSM Classification Results

Figure 2 shows the GMM-Demux classification results where each sample combination is given a distinct color. Notice that when the number of classes is large (e.g., the PBMC-2 sample), the color difference between classes become less noticeable to a naked eye.

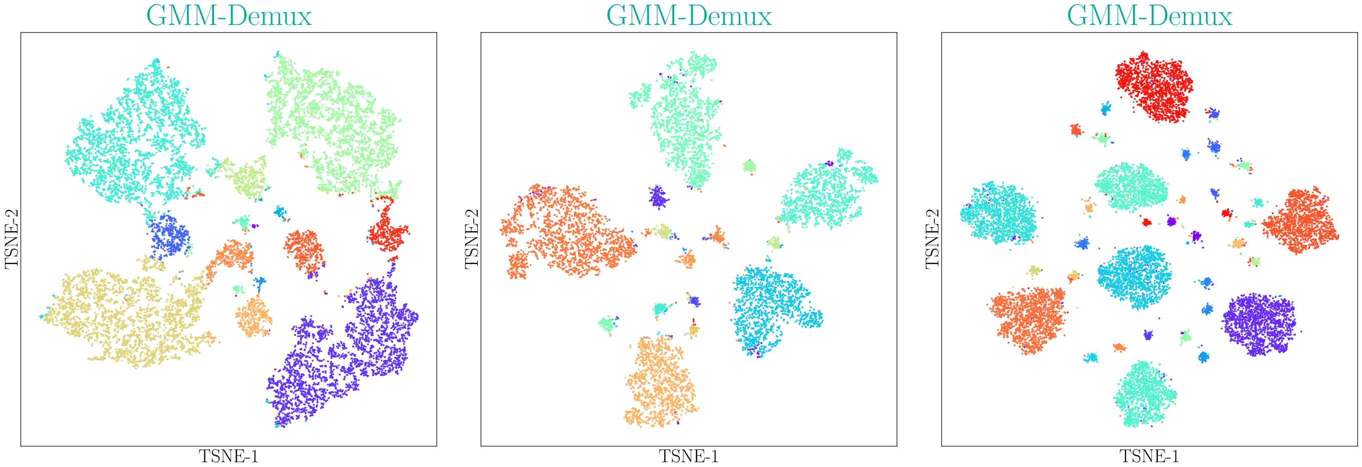


(a) PBMC-1 (b) $CD4^+$ Memory T (c) PBMC-2

Figure 2: GMM-Demux detailed classification results. Each sample and each multi-sample combination is given a unique color. As the number of classes grows, difference between colors becomes less noticeable.

### 6 Confidence Score Distributions

Figure 3 shows the confidence score distributions of the three cell hashing datasets presented in Table 2. As shown in the figure. Most droplets ($> 95\%$) have confidence scores above 0.8.

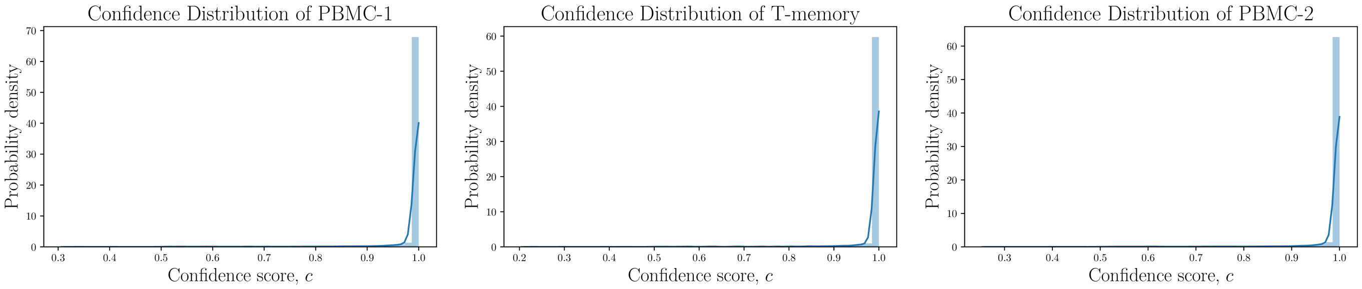


Figure 3: Confidence Score Distributions.

### 7 MSM Estimation Result for PBMC-2

The comparison between observed and model-derived MSM counts for the PBMC-2 dataset is summarized in Table 8 (provided separately). Similar to the PBMC-1 and the $CD4^+$ Memory T datasets, the observed counts and model-derived counts are highly consistent for each and every SSD and MSM category.

### 8 Sample Barcoding Profiles

This section presents sample barcoding multiplet rate profiles with number of cell-assay droplets ($X$) other than 80K.

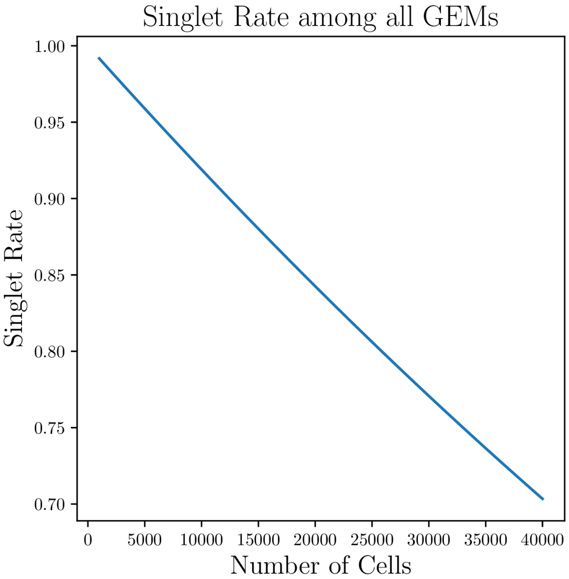


(a) Singlet rate profile.

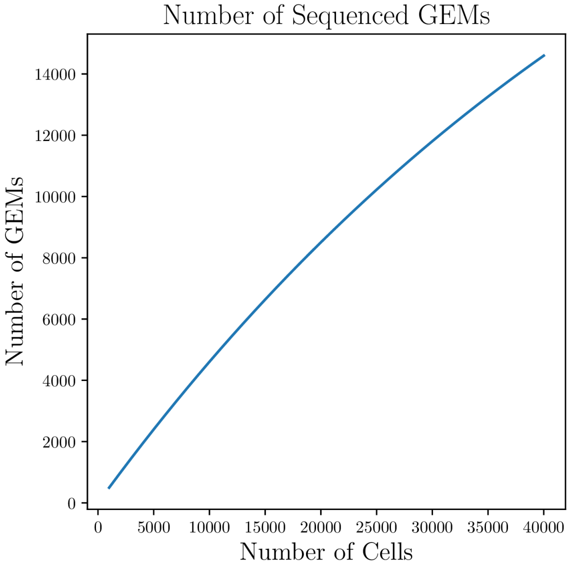


(b) GEM count profile.

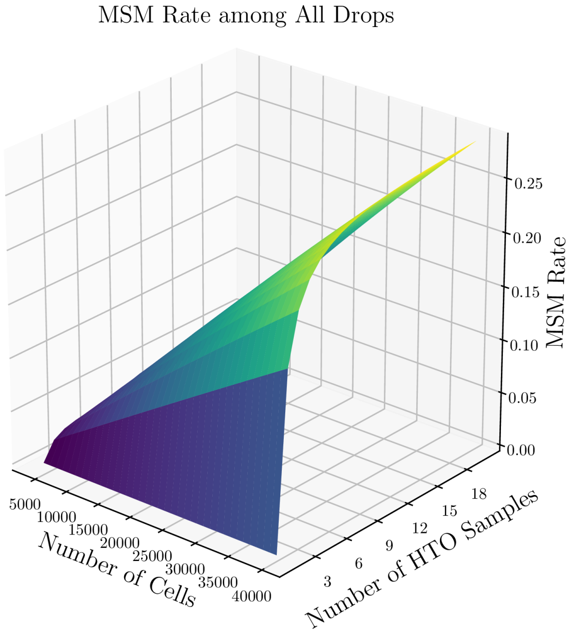


(c) MSM rate profile.

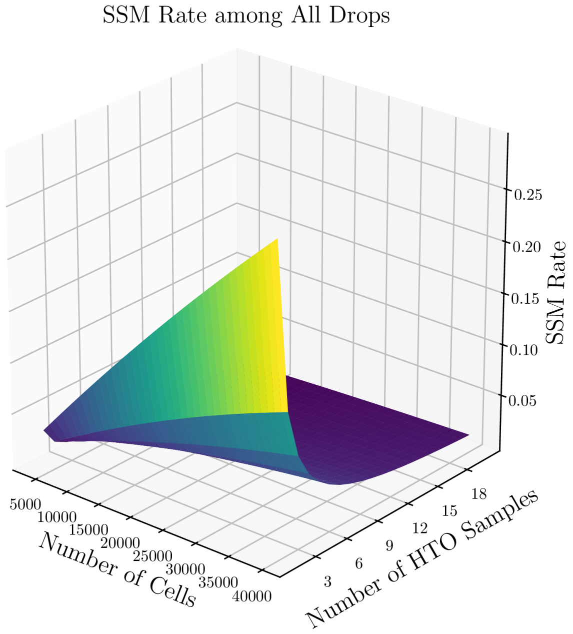


(d) SSM rate profile.

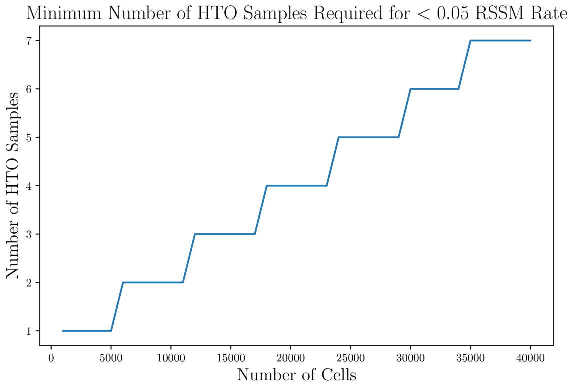


(e) Sample number profile for 5% RSSM rate target.

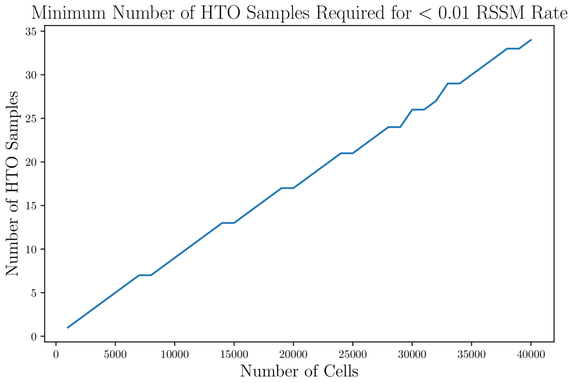


(f) Sample number profile for 1% RSSM rate.

Figure 4: Sample barcoding profiles under the assumption of 60K cell-assay droplets ($X = 60K$) and 0.5 capture rate ($r_{cap} = 50\%$).

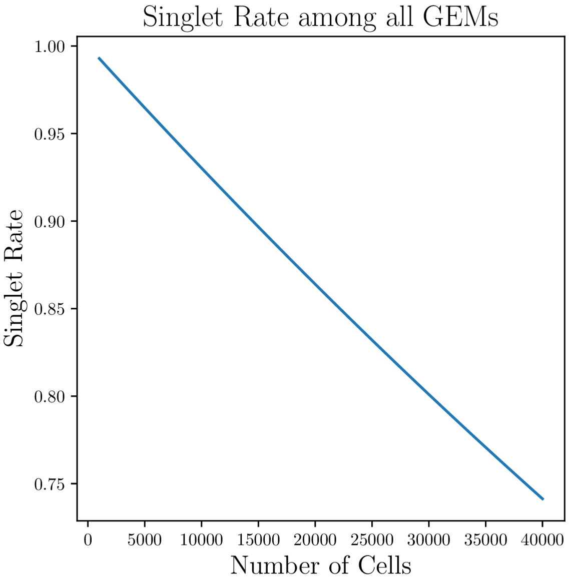


(a) Singlet rate profile.

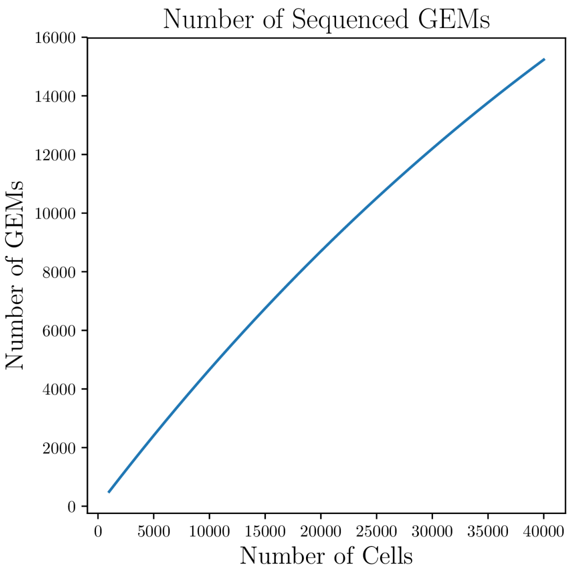


(b) GEM count profile.

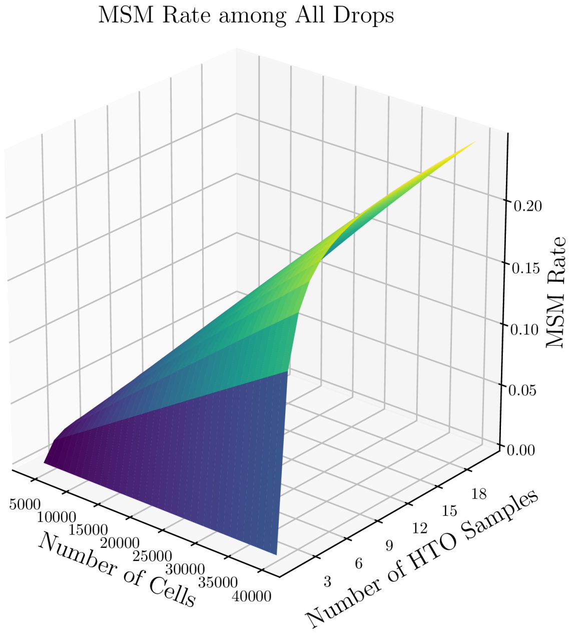


(c) MSM rate profile.

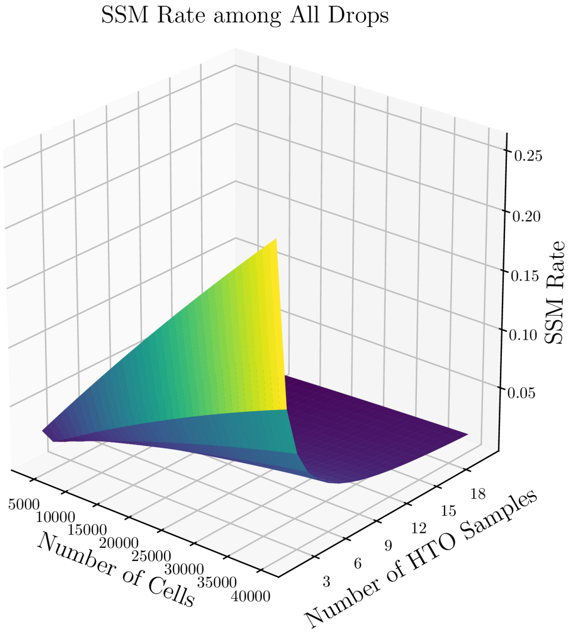


(d) SSM rate profile.

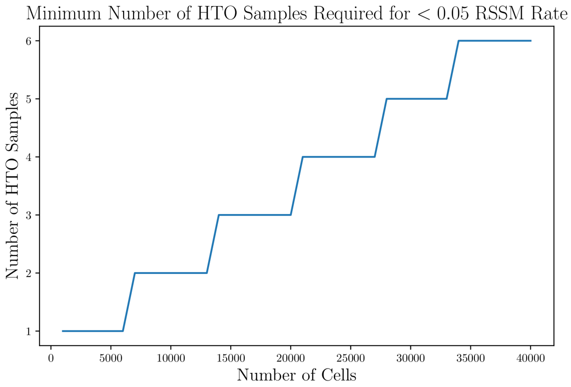


(e) Sample number profile for 5% RSSM rate target.

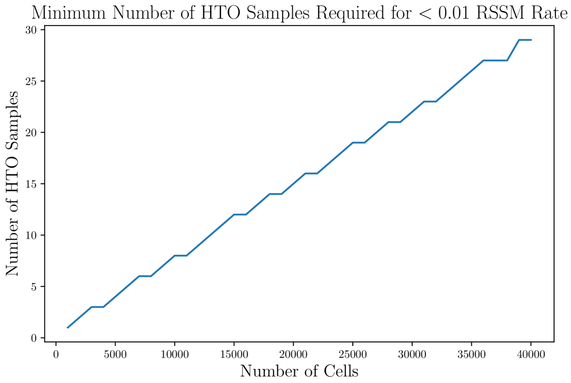


(f) Sample number profile for 1% RSSM rate.

Figure 5: Sample barcoding profiles under the assumption of 70K cell-assay droplets ($X = 70K$) and 0.5 capture rate ($r_{cap} = 50\%$).

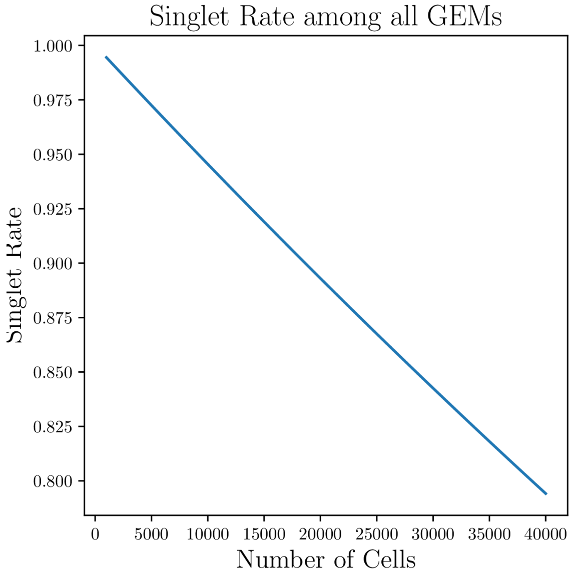


(a) Singlet rate profile.

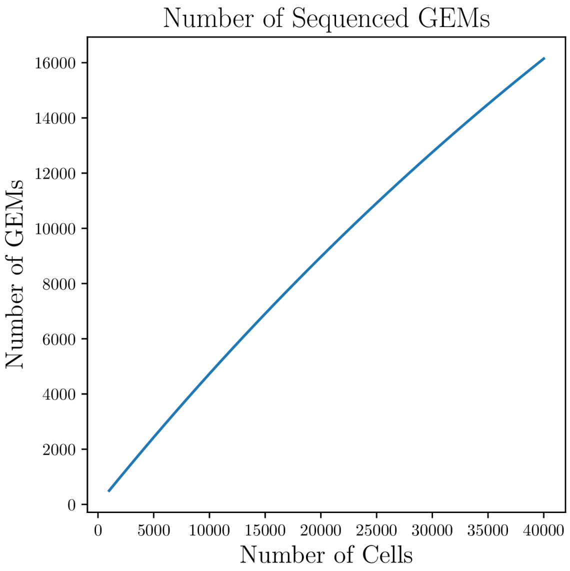


(b) GEM count profile.

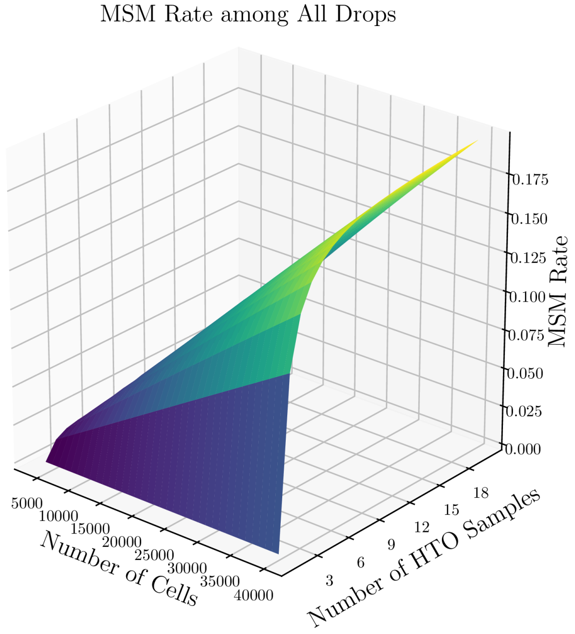


(c) MSM rate profile.

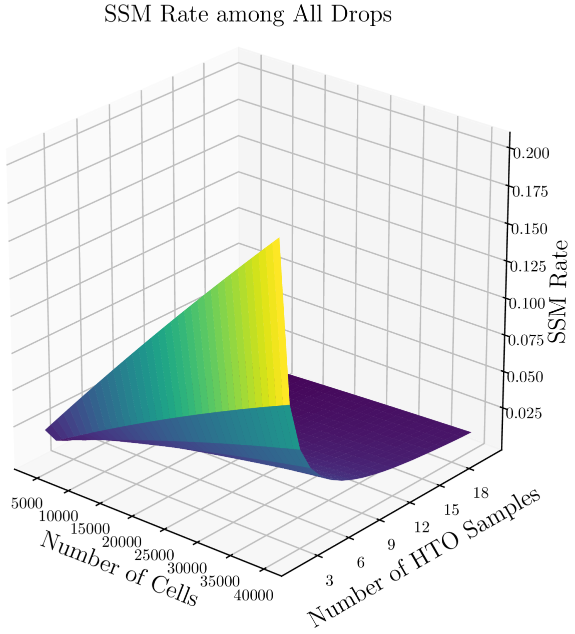


(d) SSM rate profile.

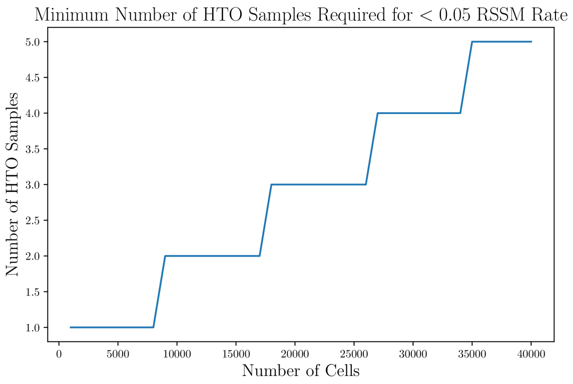


(e) Sample number profile for 5% RSSM rate target.

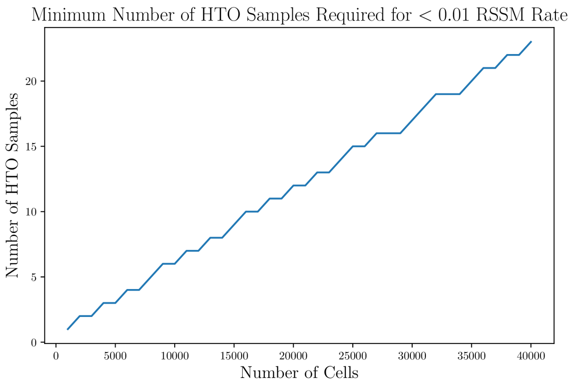


(f) Sample number profile for 1% RSSM rate.

Figure 6: Sample barcoding profiles under the assumption of 90K cell-assay droplets ($X = 90K$) and 0.5 capture rate ($r_{cap} = 50\%$).

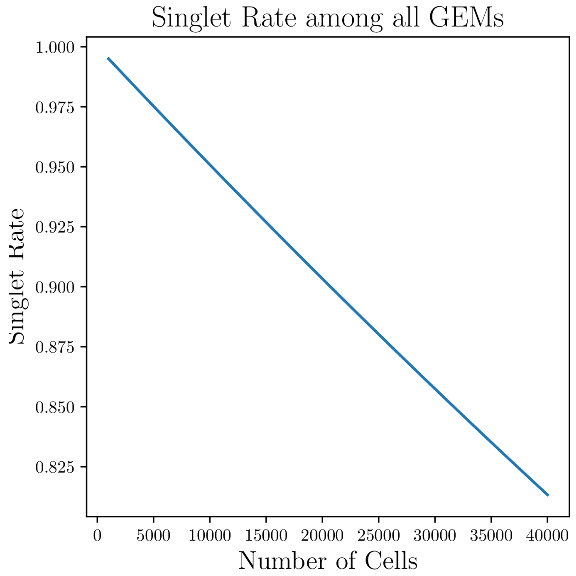


(a) Singlet rate profile.

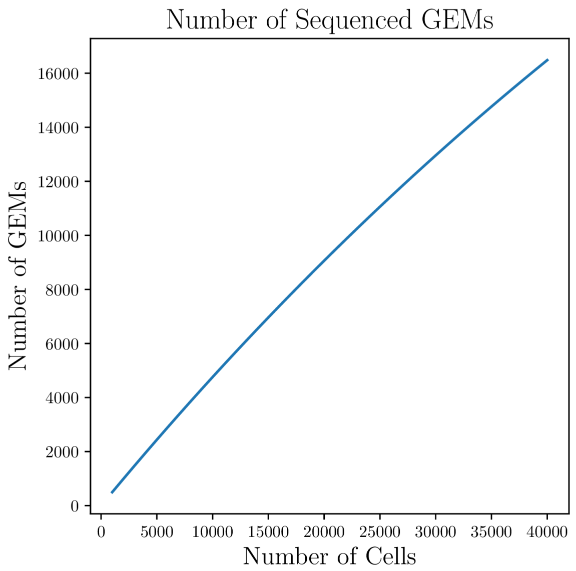


(b) GEM count profile.

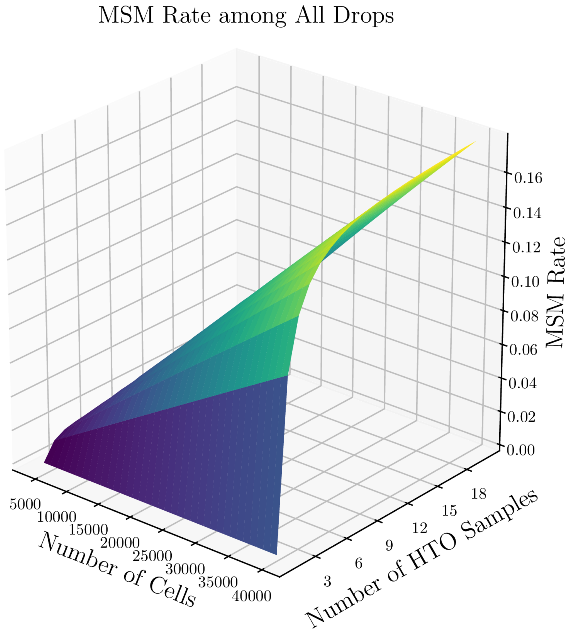


(c) MSM rate profile.

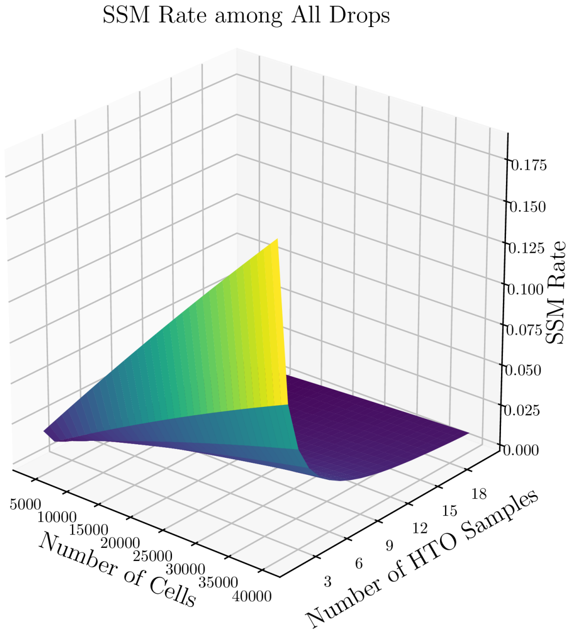


(d) SSM rate profile.

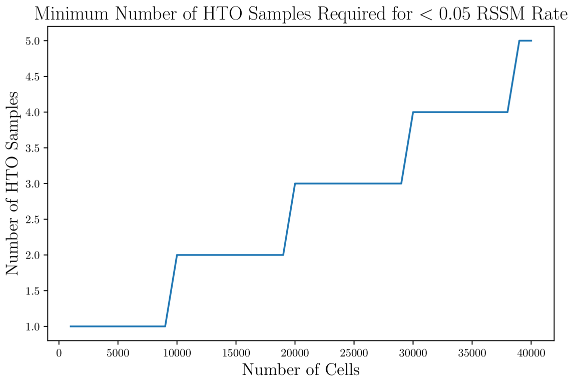


(e) Sample number profile for 5% RSSM rate target.

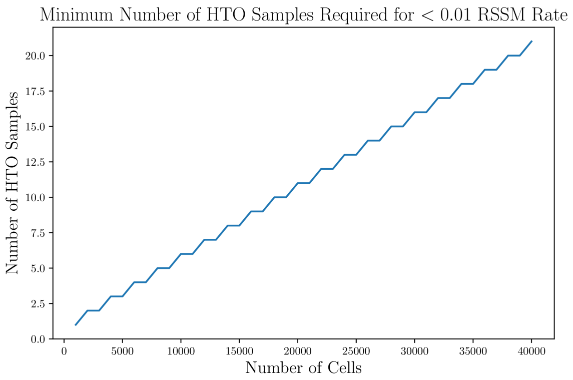


(f) Sample number profile for 1% RSSM rate.

Figure 7: Sample barcoding profiles under the assumption of 100K cell-assay droplets ($X = 100K$) and 0.5 capture rate ($r_{cap} = 50\%$).

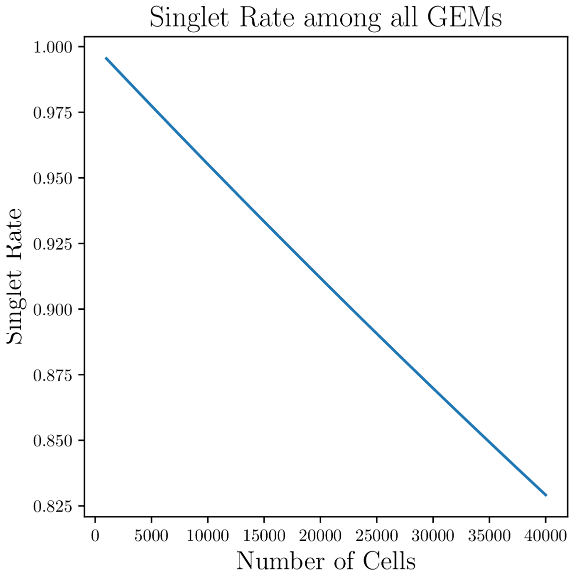


(a) Singlet rate profile.

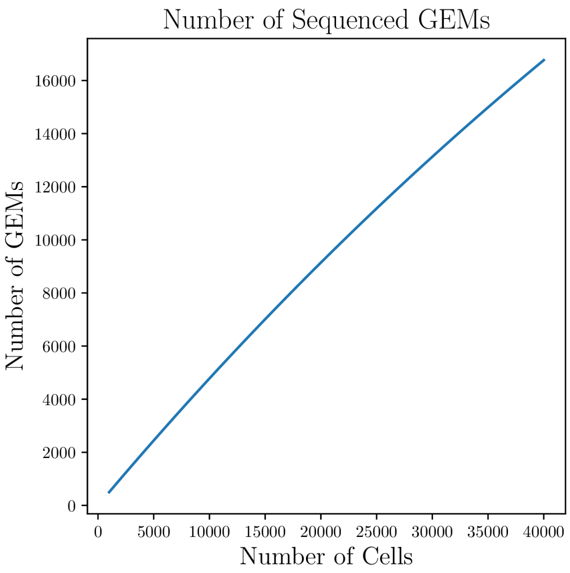


(b) GEM count profile.

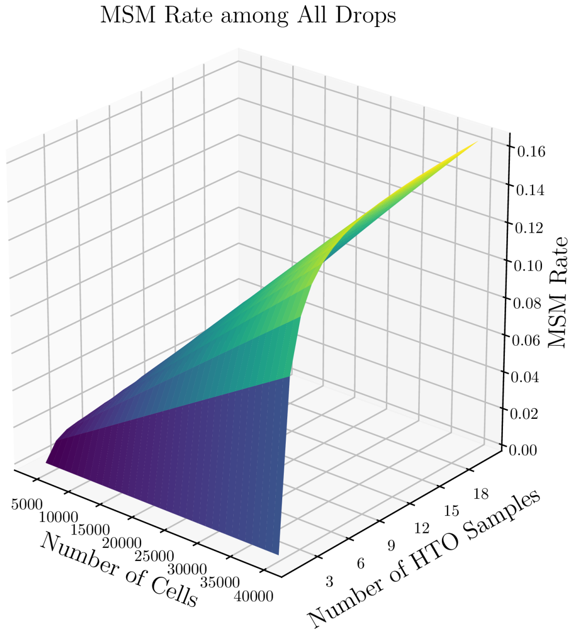


(c) MSM rate profile.

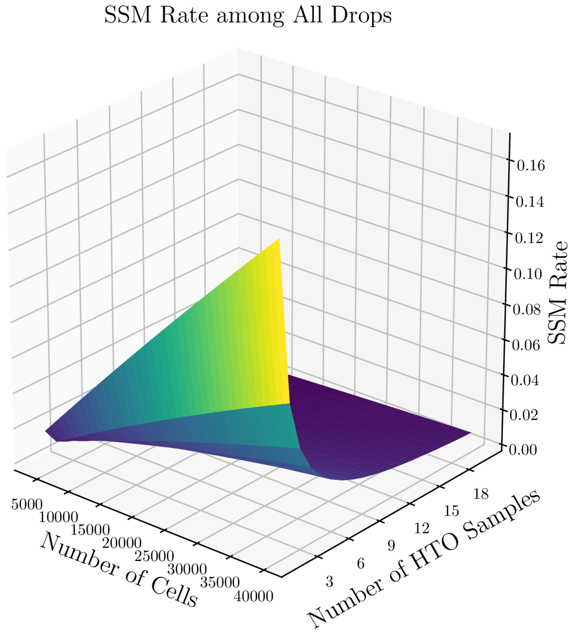


(d) SSM rate profile.

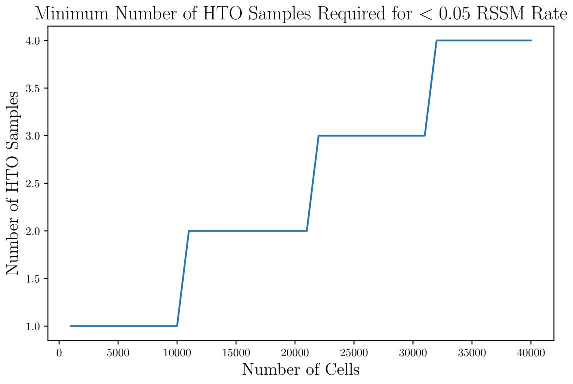


(e) Sample number profile for 5% RSSM rate target.

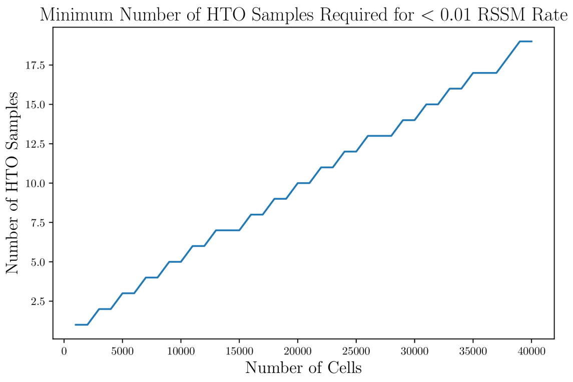


(f) Sample number profile for 1% RSSM rate.

Figure 8: Sample barcoding profiles under the assumption of 110K cell-assay droplets ($X = 110K$) and 0.5 capture rate ($r_{cap} = 50\%$).

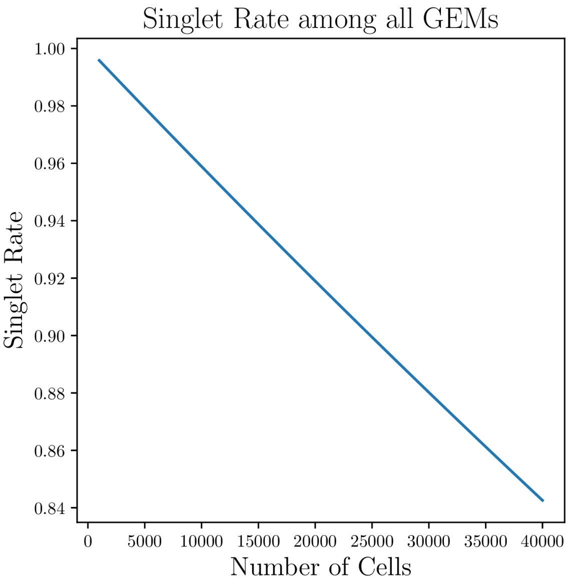


(a) Singlet rate profile.

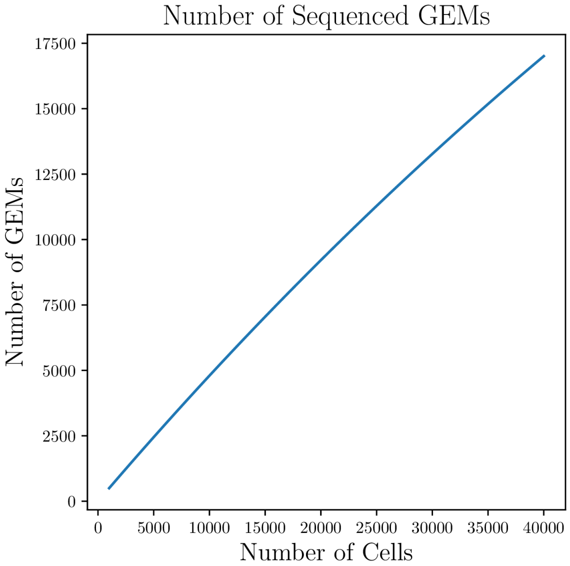


(b) GEM count profile.

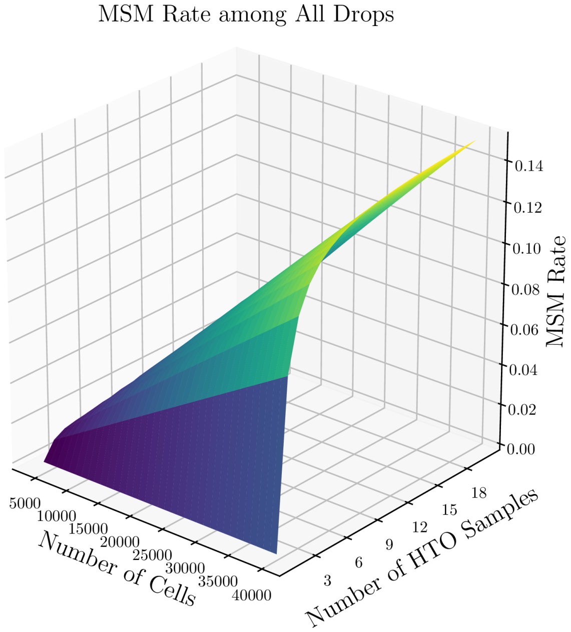


(c) MSM rate profile.

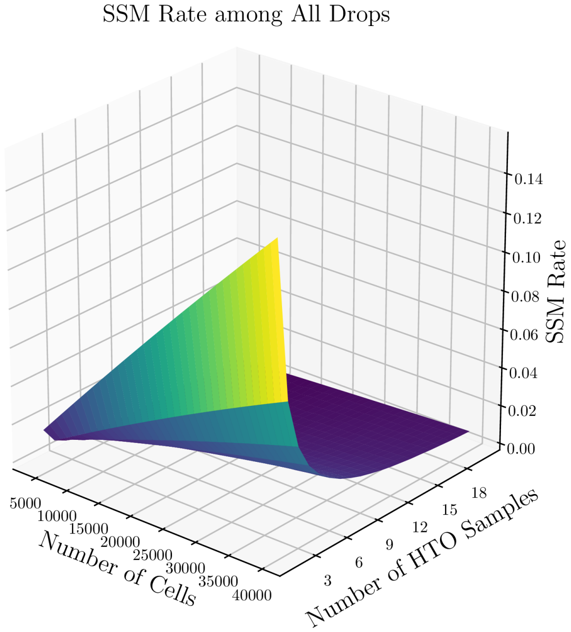


(d) SSM rate profile.

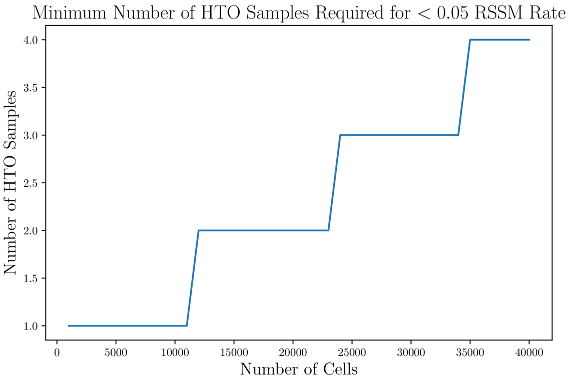


(e) Sample number profile for 5% RSSM rate target.

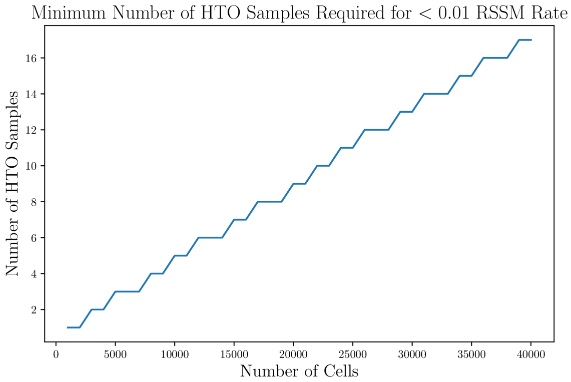


(f) Sample number profile for 1% RSSM rate.

Figure 9: Sample barcoding profiles under the assumption of 120K cell-assay droplets ($X = 120K$) and 0.5 capture rate ($r_{cap} = 50\%$).
